# Genetic or pharmacological inactivation of CREBBP sensitizes B-cell Acute Lymphoblastic Leukemia to Ferroptotic Cell Death upon BCL2 Inhibition

**DOI:** 10.1101/2023.10.02.556536

**Authors:** Alicia Garcia-Gimenez, Jonathan E. Ditcham, Dhoyazan M.A. Azazi, Eshwar Meduri, Ryan Asby, Nathalie Sakakini, Cecile K. Lopez, Nisha Narayan, Tumas Beinortas, Jaana Bagri, Shuchi Agrawal Singh, George Giotopoulos, Michael P. Murphy, Sarah J. Horton, Brian J.P. Huntly, Simon E. Richardson

**Affiliations:** Department of Haematology, Wellcome Trust—Medical Research Council Cambridge Stem Cell Institute, Cambridge, UK; Wellcome Trust—Medical Research Council Cambridge Stem Cell Institute, Cambridge, UK; Cambridge University Hospitals, Cambridge, UK; MRC Mitochondrial Biology Unit, Keith Peters Building, University of Cambridge, Cambridge, UK

## Abstract

B-cell acute lymphoblastic leukemia (B-ALL) is a leading cause of death in childhood and outcomes in adults remain dismal. There is therefore an urgent clinical need for therapies that target the highest risk cases. Mutations in the histone acetyltransferase *CREBBP* associate with high-risk features in B-ALL and have been implicated in chemoresistance. We performed a targeted drug screen in isogenic human cell lines, identifying a number of actionable small molecules that specifically target *CREBBP*-mutated B-ALL. The most potent was the BCL2 inhibitor Venetoclax, which acts through a non-canonical mechanism resulting in ferroptotic cell death. *CREBBP*-mutated cell lines showed differences in cell-cycle, metabolism and response to oxidative stress. Lastly, we demonstrate that small-molecule inhibition of CREBBP sensitizes B-ALL cells, regardless of genotype, to Venetoclax-induced ferroptosis *in-vitro* and *in-vivo*, providing a potential novel drug combination for broader clinical translation in B-ALL.

## Introduction

B-cell acute lymphoblastic leukemia (B-ALL) is an aggressive hematological malignancy of B-lineage progenitors and is the commonest cancer in children^1^. Whilst the majority of children can be cured with multi-agent chemotherapy, patients with high-risk genetic subtypes, certain age groups and those who relapse remain a clinical challenge, such that B-ALL remains a leading cause of death in childhood. Furthermore, outcomes of adults with B-ALL remain dismal, even when fit enough to be treated intensively. There is therefore a need to better understand drivers of high-risk B-ALL and to develop novel therapeutic approaches targeting these challenging patient cohorts.

*CREBBP* mutations are found in multiple hematological and solid malignancies, notably B-cell lymphomas^2,3^. Loss-of-function (LOF) mutations affecting *CREBBP* are recurrent second-hit mutations across multiple genetic subtypes of B-ALL and are associated with adverse features, including high-risk genetic subtypes and persistent measurable residual disease^4–7^. In addition, they have been mechanistically associated with chemoresistance and are enriched at relapse^4,8–12^. *CREBBP* mutations have also been described as an adverse prognostic factor in ALL, acute myeloid leukaemia (AML) and follicular lymphoma^13–15^. CREBBP is a large protein with histone acetyltransferase (HAT) enzymatic activity alongside protein scaffolding function mediated through multiple protein-protein interaction domains, including a bromodomain responsible for binding acetylated lysine residues. Alongside its paralogue EP300, CREBBP is considered to primarily function as a transcriptional co-activator, responsible for acetylating histone residues at gene enhancers and promoters. *CREBBP* LOF mutations can include complete loss of the protein or recurrent point mutations affecting the HAT domain, which appear to exert a stronger phenotype^4^. During B-ALL evolution, *CREBBP* mutations frequently become bi-allelic and commonly co-associate with activating RAS pathway mutations, suggesting strong oncogenic co-operativity^4,12,16^.

Targeting cells harbouring LOF mutations in tumour suppressor genes (TSG) principally relies on perturbing “synthetic lethal” dependencies acquired upon loss of TSG activity, commonly via inhibition of redundant pathways or protein paralogues. In the context of *CREBBP*, this has been demonstrated in models of B-cell lymphoma through inhibition of residual EP300 function using small molecule HAT or bromodomain inhibitors^17^. Global analyses of genetic co-dependencies have also implicated a dependency of *CREBBP*-mutated tumours on *EXOC5* function^18^, whilst a number of mechanistic studies have identified potentially targetable roles for CREBBP in modulating key cellular processes including DNA damage response, signaling, apoptosis and metabolism^3,12,16,19^.

In this study we identify novel treatment options for *CREBBP*-mutated high-risk B-ALL. We generated isogenic human B-ALL cell lines and undertook a synthetic lethal drug screen focusing on clinically-actionable agents targeting pathways mechanistically associated with CREBBP function. CREBBP LOF resulted in cell cycle and metabolic dysregulation associated with marked sensitivity to ferroptotic cell death upon small molecule inhibition of the anti-apoptotic regulator BCL2. Inhibition of CREBBP function with small molecule inhibitors could phenocopy this synthetic lethal effect, sensitizing diverse subtypes of B-ALL to BCL2 inhibitors *in-vitro* and producing a significant survival advantage *in-vivo*, thus providing a potential novel efficacious drug combination across a wider number of B-ALL genotypes.

## Results

### *CREBBP*-mutated B-ALL shows increased sensitivity to Venetoclax

To identify candidate therapeutics that specifically target *CREBBP*-mutated high-risk B-ALL, we undertook a synthetic-lethal drug screen. The *CREBBP* wild-type (WT) B-ALL cell line 697 (driven by the recurrent *E2A::PBX1* fusion) was genome-engineered by CRISPR-Cas9 homologous recombination to introduce a recurrent “hotspot” mutation at arginine 1446 (*CREBBP^R^*^1446^*^C^*), which is implicated as exerting a dominant-negative effect on CREBBP acetyltransferase activity^4^. Several clones were generated, including a homozygous *CREBBP^R^*^1446^*^C^* knock-in mutant clone (hereafter, 697*^KI^*) and a mutant clone containing two frameshift mutations resulting in a complete knockout of CREBBP protein (hereafter, 697*^KO^*) (Fig. 1a and Extended Data Fig. 1a). For use in validation studies, the *ETV6::RUNX1*-driven cell line REH (containing three WT copies of *CREBBP*) was also edited, resulting in two compound-heterozygous mutated clones, each including a single allele of the *CREBBP^R^*^1446^*^C^*HAT mutation, alongside presumed deleterious mutations of the other two alleles (Extended Data Fig. 1a).

**Figure 1:**
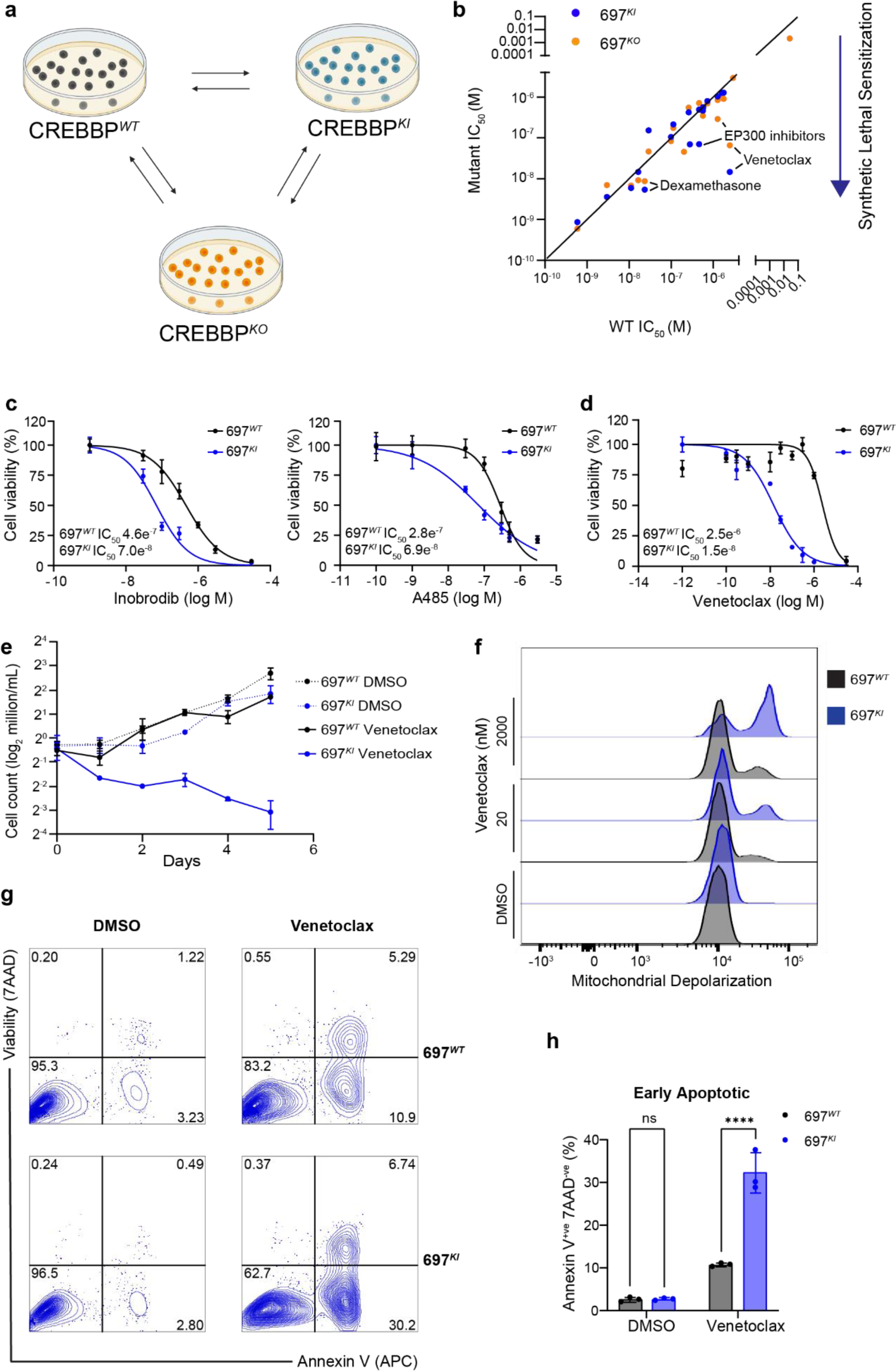
*CREBBP*-mutated B-ALL shows increased sensitivity to Venetoclax. **a,** Schematic of genome editing strategy used to engineer 697*^WT^* (black) into isogenic *CREBBP* 697*^KI^*(blue) and 697*^KO^* (yellow) clones. **b,** Cell-based drug screening was performed using a panel of 32 small molecule compounds predicted to have differential sensitivity between *CREBBP* WT and mutant lines. 72h viability was measured by MTS assays using a wide concentration range between 3pM to 30µM in triplicate and repeated using narrower concentration ranges to define IC_50_ where appropriate. Results are presented as an IC_50_ ratio of 697*^KI^* (blue) and 697*^KO^*(yellow) clones compared to WT. **c,** Dose response curves of two CREBBP/EP300 inhibitors Inobrodib (left) and A485 (right) showing enhanced sensitivity of 697*^KI^* (blue) compared to 697*^WT^* (black) in 72h MTS viability assay. Performed in triplicate, mean ± SD. **d,** Dose response curve of 697*^WT^* (black) and 697*^KI^* (blue) lines to Venetoclax in 72h MTS viability assay. Performed in triplicate, mean ± SD. **e,** Growth curve of 697*^WT^* (black) and 697*^KI^* (blue) grown in the presence of either DMSO vehicle (dotted lines) or 20nM Venetoclax (solid lines). **f,** Mitochondrial depolarization as assessed by staining for JC1 by flow cytometry (488nm 530/30) in response to DMSO vehicle, or Venetoclax at 20 or 2000nM in 697*^WT^* (black) and 697*^KI^* (blue) cell lines. **g,** Representative flow cytometry plots of externalization of Annexin-V (reported by APC) in response to 24h exposure to DMSO vehicle (left) or Venetoclax 20nM (right) in 697*^WT^* (top) and 697*^KI^*(bottom) cell lines. Viability is assessed by 7AAD exclusion. **h,** Summary of 3 replicate experiments measuring proportion of viable 7AAD^-ve^Annexin-V^+ve^ early apoptotic cells. Mean ± SD, 2 way ANOVA ****, *P* <0.0001.

We subjected the 697 isogenic cell lines to a targeted drug screen, using a wide range of concentrations, focussed on clinically-actionable drugs in classes implicated or hypothesized to show differential sensitivity in published models of B-cell lymphoma and other *CREBBP*-mutated malignancies (Extended Data Table 1)^3,4,10–12,17,19,20^. *CREBBP*-mutated 697 cells were not differentially sensitive to traditional cytotoxic chemotherapy, and paradoxically showed a degree of sensitization to the glucocorticoid Dexamethasone, used in current ALL induction regimens (Fig. 1b and Extended Data Table 1 and Fig. 1b)^4,11,12^. As anticipated, and validating our screen design, inhibitors of the CREBBP paralogue EP300 (the CREBBP/EP300-specific bromodomain inhibitor Inobrodib and the CREBBP/EP300 acetylase inhibitor A485) exhibited synthetic lethality, consistent with previous reports in B-cell lymphoma (Fig. 1b,c and Extended Data Table 1 and Fig. 1c)^17^.

Unexpectedly, the most potent hit identified from the screen was the clinical-grade BCL2 inhibitor Venetoclax, which showed a 2-log_10_-fold reduction in IC_50_, in both 697*^KI^* and 697*^KO^* clones (Fig. 1b,d and Extended Data Table 1 and Fig. 1d). These findings were validated in isogenic REH lines, with a 1-log_10_-fold reduction in IC_50_ in both mutant clones (Extended Data Fig. 1e). We confirmed this sensitization to Venetoclax in *in-vitro* proliferation assays by direct cell counting (Fig. 1e). Upon low-dose Venetoclax exposure, *CREBBP*-mutated 697 cells showed enhanced evidence of markers of programmed cell death, including mitochondrial depolarization, externalization of Annexin-V and induction of cleaved PARP and caspase-3 (Fig.1 f-h and Extended Data Fig. 1f), consistent with the known mechanism-of-action of Venetoclax in inducing apoptotic programmed cell death.

Overall, this focussed drug screen demonstrates that *CREBBP*-mutated B-ALL is: i) not uniformly chemo-resistant, and ii) identifies a number of clinically-actionable agents for use in *CREBBP*-mutated high-risk B-ALL, including Dexamethasone, EP300 inhibitors and a potent sensitization to the BCL2 inhibitor Venetoclax.

### Venetoclax exerts its effect on *CREBBP*-mutated B-ALL by on-target inhibition of BCL2

We sought to explore the mechanism-of-action of Venetoclax. Venetoclax was developed to induce apoptosis through inhibition of BCL2 binding to the pro-cell death proteins BAK and BAX. However, recently, it has been shown to have alternative mechanisms-of-action, in particular on metabolism and self-renewal, including potential BCL2-independent effects^21–23^. To test whether the mechanism-of-action of Venetoclax in *CREBBP*-mutated B-ALL was through on-target BCL2 inhibition, we employed a doxycycline-inducible shRNA knock-down system, where shRNA expression was directly linked to a fluorescent reporter (Extended Data Fig. 2a-d)^24^.

Co-culture of 697*^WT^* or 697*^KI^* cells expressing one of two unique *BCL2*-targeting shRNAs (reported by mCherry) competed against cells expressing an shRNA targeting the negative-control *renilla* gene (reported by green fluorescent protein (GFP)) showed that 697*^KI^* cells exhibited a marked competitive disadvantage upon *BCL2*-knockdown, when compared to 697*^WT^* (Fig. 2a,b). This was associated with significant externalization of Annexin-V, consistent with the induction of programmed cell death (Fig. 2c,d).

**Figure 2:**
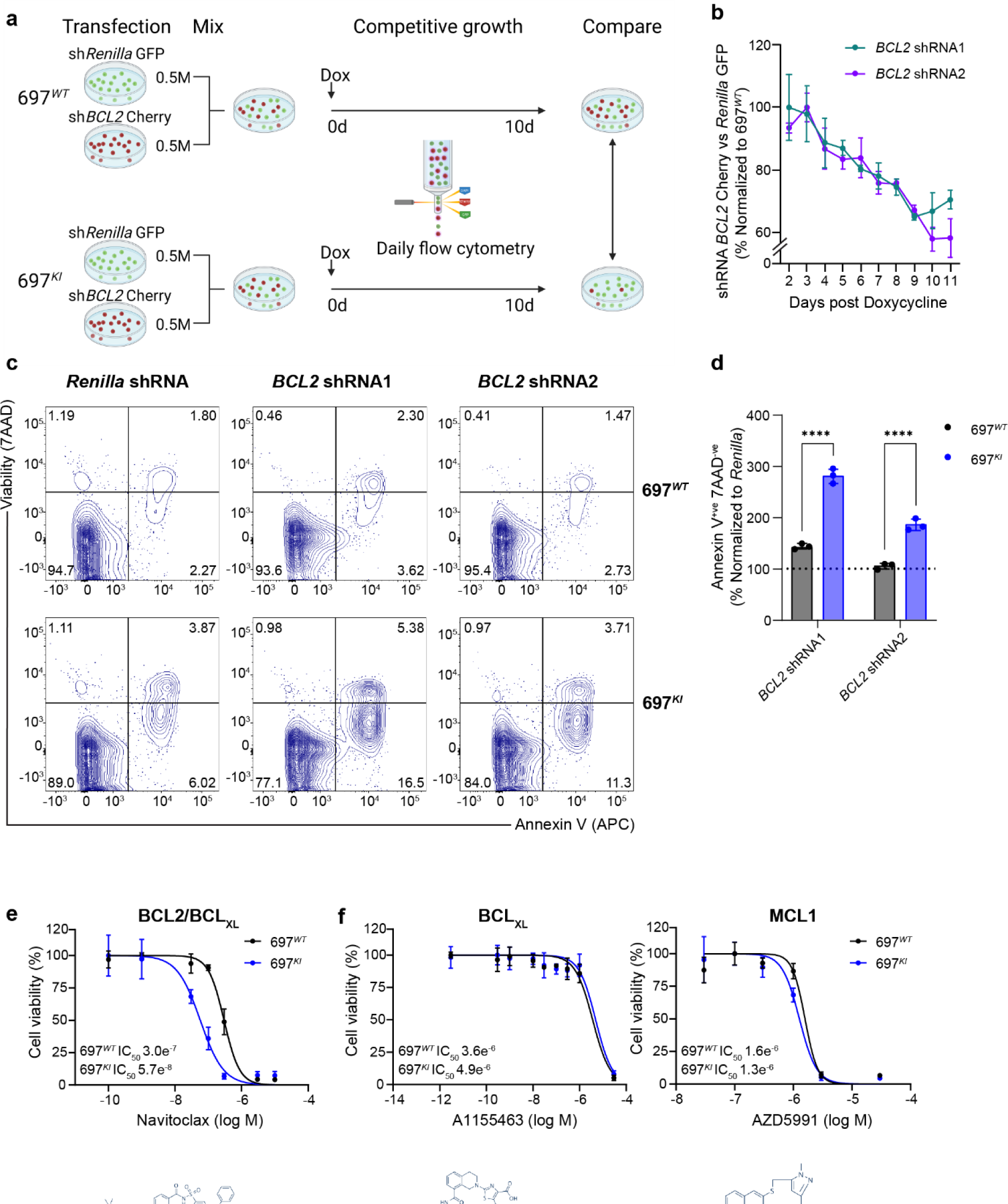
Venetoclax exerts its effect on *CREBBP*-mutated B-ALL by on-target inhibition of BCL2. **a,** Schematic of doxycycline-inducible shRNA KD system competitive co-culture assay. 697*^WT^* and 697*^KI^* cells were stably transfected with two separate doxycycline-inducible shRNAs targeting *BCL2,* reported by mCherry, or a control shRNA targeting *Renilla*, reported by GFP. *BCL2* and *Renilla* shRNA-expressing cells were mixed in equal numbers, doxycycline added to the media (500ng/ml) to induce shRNA expression and the proportion of cells surviving *BCL2/Renilla*-KD analysed by daily flow cytometry. Created with BioRender.com. **b,** *BCL2* shRNA KD competitive proliferation assay for two different *BCL2*-targeting shRNAs are presented, showing the ratio of *BCL2*-targeting shRNA (mCherry) vs. *Renilla* control (GFP), in 697*^KI^* normalized to 697*^WT^* as a percentage. Performed in triplicate, mean ± SD. **c,** Representative flow cytometry plots of externalization of Annexin-V (APC) in 697*^WT^* (top) and 697*^KI^* (bottom) cell lines in response to doxycycline-induced shRNAs targeting *Renilla* (left), or two different *BCL2-*targeting shRNAs (middle and right). Day 6 post induction. Viability is assessed by 7AAD exclusion. **d,** Summary of 3 replicate experiments measuring proportion of viable 7AAD^-ve^Annexin-V^+ve^ early apoptotic cells normalized to doxycycline-induced *Renilla* control. Analysed 6 days after induction, performed in triplicate, mean ± SD, 2 way ANOVA ****, *P* <0.0001. **e,** Dose response curve of 697*^WT^* (black) and 697*^KI^*(blue) to Navitoclax in 72h MTS viability assays. Performed in triplicate, mean ± SD. Chemical structure of Navitoclax is shown below. Images from Selleckchem. **f,** Dose response curve of 697*^WT^*(black) and 697*^KI^* (blue) to A1155463 (BCL_XL_ only inhibitor, left) and AZD5991 (MCL1 inhibitor, right) in 72h MTS viability assays. Performed in triplicate, mean ± SD. Chemical structure of drugs are shown below. Images from Selleckchem.

Furthermore, 697*^KI^* cells also showed differential sensitivity to the structurally-unrelated clinical-grade dual BCL2/BCL_XL_ inhibitor Navitoclax (Fig. 2e and Extended Data Fig. 2e,f). Conversely, specific inhibitors of other antiapoptotic proteins (BCL_XL_ and MCL1) showed no specificity for *CREBBP*-mutated cells (Fig. 2f), confirming a BCL2-specific effect.

Collectively these studies demonstrate that Venetoclax induces programmed cell death in *CREBBP*-mutated B-ALL by on-target BCL2 inhibition and that the sensitivity of 697*^KI^*cells is specific to BCL2 and does not occur through other anti-apoptotic proteins.

### *CREBBP*-mutated B-ALL shows significant cell cycle and metabolic dysregulation

To further explore the mechanism of action of Venetoclax in *CREBBP*-mutated 697 cells, we undertook bulk RNA sequencing (RNAseq) of 697*^WT^* and 697*^KI^*cells, after 24 hours exposure to either dimethyl sulfoxide (DMSO) vehicle, or low-dose Venetoclax (20nM – the IC_50_ of 697*^KI^* cells). Consistent with the role of *CREBBP* as a transcriptional co-activator, the majority of differentially expressed genes (DEGs) between DMSO-vehicle treated 697*^WT^* and 697*^KI^* cells were down-regulated (Extended Data Fig.3a). Gene Set Enrichment Analysis (GSEA) showed marked down-regulation of published *Crebbp* target genes from a mouse lymphoma model, supporting the role of the *CREBBP^R^*^1446^*^C^* mutation as inhibitory to CREBBP transcriptional co-activator function (Extended Data Fig. 3b)^25^.

KEGG pathway and GSEA analysis showed significant down-regulation of signatures associated with apoptosis, in consonance with our functional experiments above (Fig. 1f-h and Fig. 3a,b). However, gene-specific examination of differential expression of apoptotic regulators by KEGG pathway analysis showed a mixed picture, affecting the expression of both pro- and anti-apoptotic genes, including a small but significant up-regulation of *BCL2* itself (Fig. 3c and Extended Data Fig. 3c).

**Figure 3:**
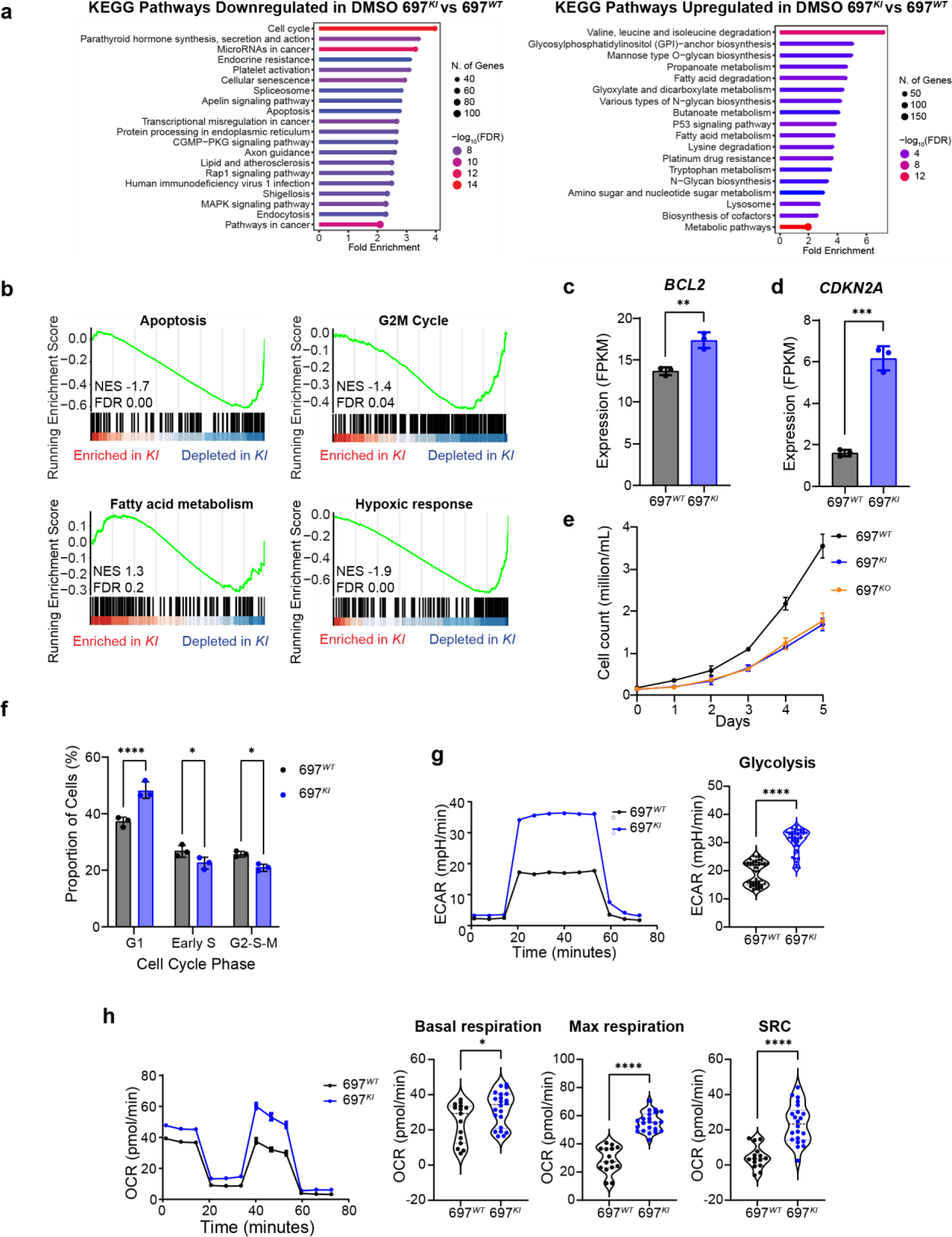
*CREBBP*-mutated B-ALL shows significant cell cycle and metabolic dysregulation. **a,** Summary of most significantly down-regulated (left) and up-regulated (right) KEGG (Kyoto Encyclopaedia of Gene and Genomes) pathways from DEGs of RNAseq analysis comparing DMSO vehicle-treated 697*^KI^* with 697*^WT^*. **b,** GSEA analysis of ranked genes of RNAseq analysis comparing DMSO vehicle-treated 697*^KI^* with 697*^WT^*. NES: normalized enrichment score; FDR: false discovery rate. **c,** Comparison of *BCL2* expression levels in 697*^WT^* (black) and 697*^KI^* (blue) cells by RNAseq. Each dot represents one sample. Bars show mean fragments per kilobase of transcript per million fragments mapped (FPKM) value ± SD, n=3, significance calculated by unpaired *t* test, **, *P*=0.0038. **d,** Comparison of *CDKN2A* expression levels in 697*^WT^* (black) and 697*^KI^* (blue) cells by RNAseq. Each dot represents one sample. Bars show mean FPKM value ± SD, n=3, significance calculated by unpaired *t* test, ***, *P*=0.0002. **e,** Proliferation of untreated 697*^WT^* (black), 697*^KI^*(blue) and 697*^KO^* (yellow) cells measured by direct counting. Performed in triplicate, mean ± SD. **f,** Analysis of cell cycle stage by FUCCI reporter system. Percentage proportion of cells in G1, Early S and G2-S-M phases in 697*^WT^* (black) and 697*^KI^* (blue) cells. N=3, each dot represents mean average of 3 technical replicates, bar shows mean average; significance calculated by 2 way ANOVA, ****, *P* <0.0001, Early S *P*=0.0498; G2-S-M *P*=0.0342. **g,** Glycolytic rate measured by extracellular acidification rate (ECAR) using Seahorse (Agilent) Glycostress test in 697*^WT^* (black) and 697*^KI^* (blue) cells. Left panel: Representative ECAR plot over time. Mean ± SEM. Right panel: Summary of maximal ECAR. Each dot represents a single replicate acquired from two separate experiments. Significance calculated by unpaired *t* test, ****, *P* <0.0001. **h,** Mitochondrial oxygen consumption rate (OCR) measured using Seahorse (Agilent) Mitostress test. Left panel: Representative OCR plot over time. Mean ± SEM. Right panel: Summary of basal OCR (left), maximal OCR (middle) and spare respiratory capacity (right). Each dot represents a single replicate acquired from two separate experiments. Significance calculated by unpaired *t* test, ****, *P* <0.0001; *, *P*=0.0315.

More broadly, KEGG and GSEA pathway analyses showed a differential down-regulation of cell cycle and signaling pathways in 697*^KI^*(Fig. 3a,b and Extended Data Fig. 3d). Downregulation of cell cycle-associated transcriptional signatures was associated with a marked up-regulation of the tumour suppressor *CDKN2A* (encoding the negative cell-cycle regulator P16^INK4a–ARF^), which is commonly mutated in B-ALL (Fig. 3d). We confirmed a relative reduction in proliferative capacity in both 697*^KI^*and 697*^KO^* cells compared to 697*^WT^* by proliferation assays (Fig. 3e). This was associated with a significantly increased proportion of 697*^KI^* cells in G1 phase alongside reduced Early S/G2-S-M phases (Fig. 3f and Extended Data Fig. 3e) confirming a significant defect in cell cycle progression. The majority of transcriptionally up-regulated KEGG and Gene Ontology (GO) pathways were indicative of metabolic dysfunction, and GSEA showed dysregulation of fatty acid oxidation and hypoxic gene signatures (Fig. 3a,b and Extended Data Fig. 3f). Given the close association of cell cycle and metabolism, and the established role of BCL2 and Venetoclax in disturbing mitochondrial respiration^23^, we analysed baseline metabolic differences between 697*^WT^* and 697*^KI^* cells using *in-vitro* metabolic flux assays. Unexpectedly, and despite lower cell cycle progression, 697*^KI^* cells consistently showed increased rates of both glycolysis and oxidative phosphorylation (OxPhos), including higher rates of both basal and maximal respiration, and an increase in spare respiratory capacity (SRC) (Fig. 3g,h).

Collectively our model suggests that loss of CREBBP acetyltransferase function results in significant transcriptional dysregulation, affecting multiple cellular processes including apoptosis, cell cycle and metabolism.

### Venetoclax induces ferroptotic cell death in *CREBBP*-mutated B-ALL

To explore the transcriptional impact of low-dose Venetoclax treatment of 697*^KI^* cells, we employed a four-way interaction model to identify genes specifically dysregulated in 697*^KI^* cells upon Venetoclax treatment (Extended Data Fig. 4a). KEGG pathway analyses of these genes showed further down-regulation of cell cycle-associated genes and enrichment for metabolic pathways and ferroptosis in Venetoclax-treated 697*^KI^* cells (Fig. 4a). Furthermore, GSEA showed marked up-regulation of genes associated with multiple metabolic processes, ROS scavenging, ferroptosis and the unfolded protein response (Fig. 4b).

**Figure 4:**
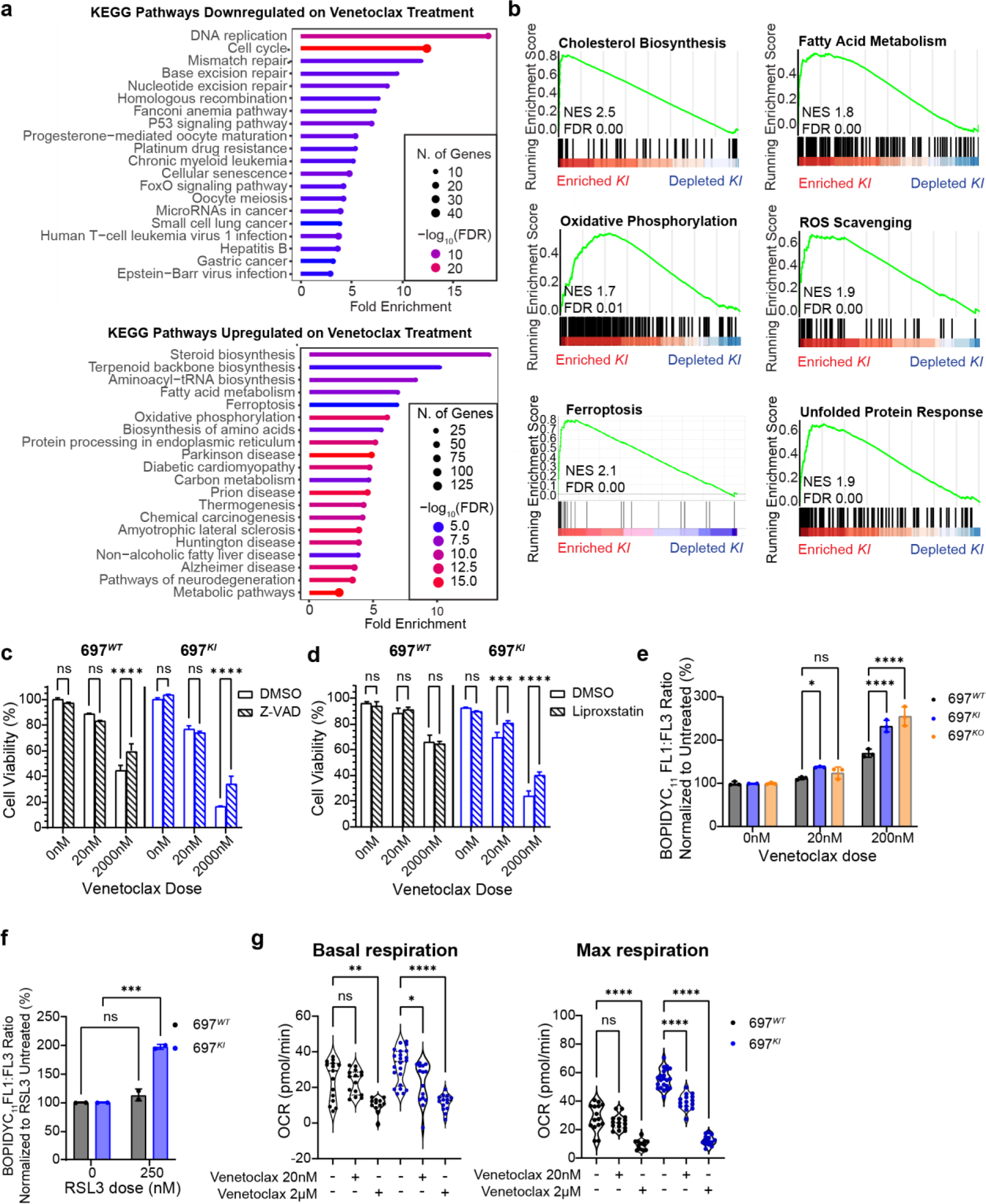
Venetoclax induces ferroptotic cell death in *CREBBP*-mutated B-ALL. **a,** Summary of most significantly down-regulated (top) and up-regulated (bottom) KEGG pathways from DEGs of RNAseq analysis comparing Venetoclax-treated 697*^KI^* with 697*^WT^*. **b,** GSEA analysis of ranked genes of RNAseq analysis comparing Venetoclax-treated 697*^KI^* with 697*^WT^*. **c,** Cellular viability assessed by flow cytometry of 697*^WT^* (black) and 697*^KI^* (blue) cells treated with increasing doses of Venetoclax at 24h either with exposure to the cell-permeant pan-caspase inhibitor Z-VAD (hashed) or DMSO control (plain). Performed in triplicate, mean ± SD, analysed by 2 way ANOVA, ****, *P* <0.0001. **d,** Cellular viability assessed by flow cytometry of 697*^WT^* (black) and 697*^KI^* (blue) cells treated with increasing doses of Venetoclax at 24h, either with exposure to the ferroptotic inhibitor Liproxstatin 1 (hashed) or DMSO control (plain). Performed in triplicate, mean ± SD, analysed by 2 way ANOVA, ****, *P* <0.0001; ***, *P*=0.0004. **e,** Lipid peroxidation in Venetoclax-treated 697*^WT^* (black), 697*^KI^* (blue) and 697*^KO^* (yellow) cells assessed by BODIPYC_11_ expressed as a ratio of FL1:FL3 (488nm 530/30:610/20) mean fluorescence intensity (MFI) normalized to untreated cells. Performed in triplicate, each dot represents a single sample, mean ± SD, two way ANOVA, ****, *P* <0.0001; *, *P*=0.0179. **f,** Lipid peroxidation in RSL3-treated 697*^WT^*(black) and 697*^KI^* (blue) cells assessed by BODIPYC_11_ expressed as a ratio of FL1:FL3 (488nm 530/30:610/20) mean fluorescence intensity (MFI) normalized to untreated cells. Mean ± SD of 2 experiments performed in triplicate. Two way ANOVA, ***, *P*=0.0001. **g,** Summary of basal (left) and maximal (right) mitochondrial oxygen consumption rate (OCR) measured using Seahorse (Agilent) Mitostress test in 697*^WT^*(black) and 697*^KI^* (blue) cells exposed to increasing concentrations of Venetoclax. Each dot represents a single replicate acquired from two separate experiments. One way ANOVA ****, *P* <0.0001; **, *P*=0.0012; *, *P*=0.0176 .

Ferroptosis is a distinct form of programmed cell death resulting from iron-catalyzed reactive oxygen species (ROS)-mediated damage to unsaturated fatty acids of membrane phospholipids^26^. It commonly co-associates with apoptosis and is associated with expression cell death markers, including Annexin-V externalization^27^. We demonstrated evidence of ferroptosis upon Venetoclax treatment specifically occurring in 697*^KI^* cells using *in-vitro* assays. Exposure to the cell-permeable pan-caspase inhibitor Z-VAD partially rescued viability to high-dose Venetoclax in both 697*^WT^* and 697*^KI^* cells, indicating a role for intrinsic, caspase-mediated apoptosis induced by high-dose Venetoclax in both lines (Fig. 4c). Conversely, exposure to Liproxstatin 1, which specifically inhibits ferroptosis-associated lipid peroxidation, rescued viability in 697*^KI^*cells exposed to either low or high-dose Venetoclax (Fig. 4d), with no effect seen in 697*^WT^*cells. BCL2 inhibition with either Venetoclax or Navitoclax was also associated with ferroptosis in 697*^KI/KO^* and REH*^Mut^*cells, as shown by elevated BODIPYC_11_ staining, an indicator of lipid peroxidation (Fig. 4e and Extended Data Fig. 4b-d).

We tested the intrinsic susceptibility of 697*^WT^* and 697*^KI^* cells to ferroptosis. The inhibition of glutathione phospholipid peroxidase activity using the GPX4 inhibitor RSL3 induced significantly higher levels of lipid peroxidation in 69*7^KI^* cells, further demonstrating their enhanced sensitivity to redox stress (Fig. 4f). Metabolically, low dose (20nM) Venetoclax specifically resulted in a reduction in both basal and maximal OxPhos in 697*^KI^* cells, whereas only high-dose Venetoclax (2000nM) resulted in a marked reduction in OxPhos in both 697*^KI^* and 697*^WT^* cells, consistent with BCL2’s role in regulating mitochondrial outer membrane permeabilization (Fig. 4g).

Overall, these findings demonstrate that the major driver underlying the sensitivity of *CREBBP*-mutated cells to BCL2 inhibition is ferroptotic programmed cell death, associated with underlying metabolic dysregulation.

### Pharmacological inhibition of CREBBP function can sensitize B-ALL to Venetoclax *in-vitro*

The majority of B-ALL patients do not have *CREBBP*-mutations. Given the strength of the association with Venetoclax sensitization we hypothesized that pharmacological inhibition of *CREBBP^WT^* function could sensitize *CREBBP^WT^*B-ALL to BCL2 inhibitors.

Pre-treatment of 697*^WT^* or REH*^WT^* cells with the preclinical CREBBP/EP300 HAT inhibitor A485 almost perfectly phenocopied the Venetoclax sensitization seen in our isogenic lines (Fig. 5a and Extended Data Fig. 5a). Furthermore, co-treatment with A485 and Venetoclax showed strong pharmacological synergy (Fig. 5b). Further corroborating our isogenic findings, A485-sensitized Venetoclax cytotoxicity was associated with enhanced lipid peroxidation consistent with ferroptosis (Fig. 5c and Extended Data Fig. 5b). Similar results were also seen with the early-phase CREBBP/EP300-specific bromodomain inhibitor Inobrodib (Extended Data Fig. 5c,d). In both 697*^WT^* and REH*^WT^*cells, A485-induced HAT inhibition was associated with increased basal and maximal OxPhos with an increase in SRC, similar to that seen in 697*^KI^* cells (Fig. 5d and Extended Data Fig. 5e-g). Single-agent A485-treated 697*^WT^* cells showed no loss of proliferation or evidence of programmed cell death by annexin-V externalization, strongly suggesting that at these doses A485 works by sensitizing cells to Venetoclax-induced cytotoxicity (Fig. 5e,f).

**Figure 5:**
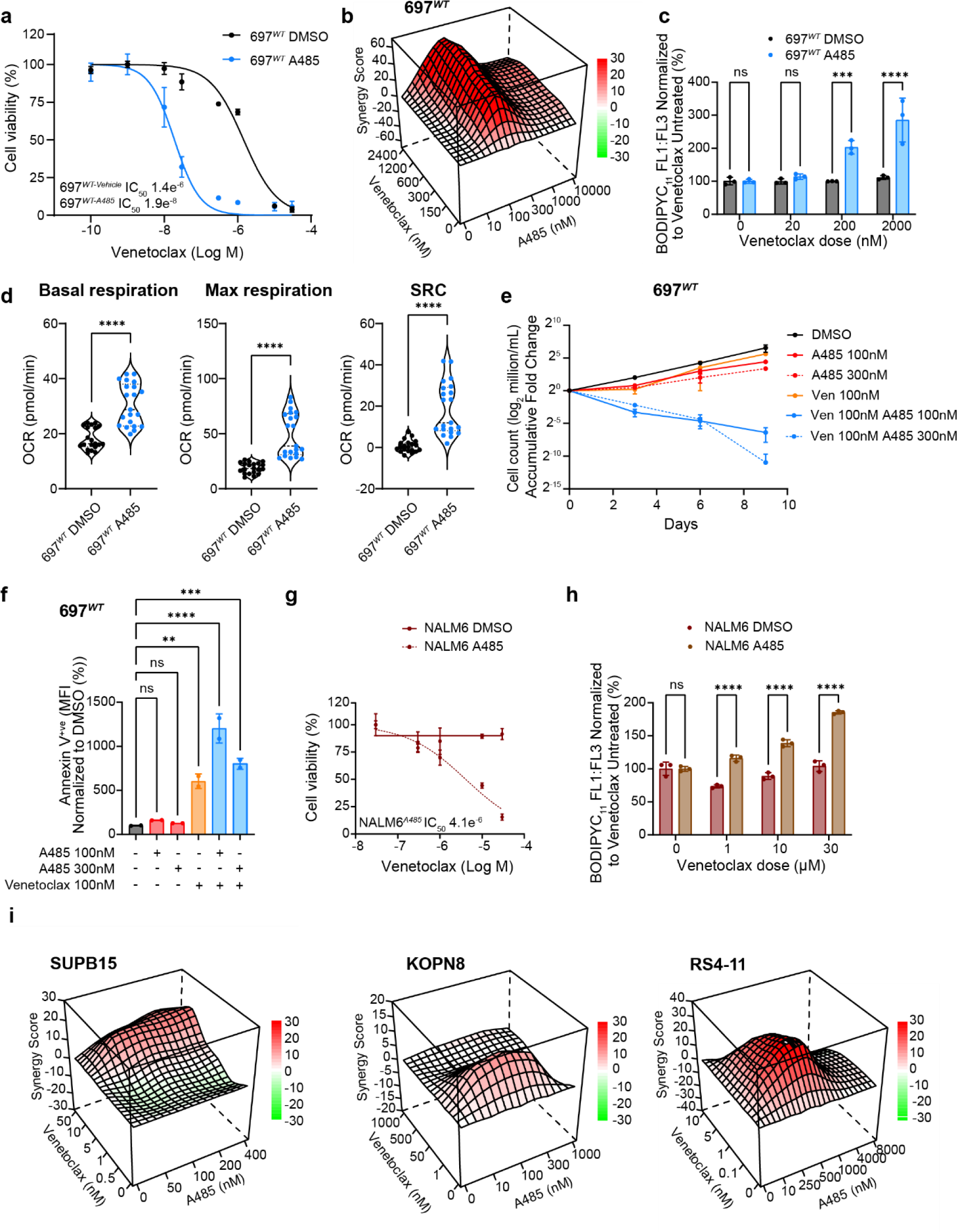
Pharmacological inhibition of CREBBP function can sensitize B-ALL to Venetoclax *in-vitro*. **a,** Dose response curve of 697*^WT^* (black) and 697*^WT^* pre-treated with 3 days of A485 (pale blue) to Venetoclax in 72h MTS viability assays. Performed in triplicate, mean ± SD. **b,** Three-dimensional diffusion plot of ZIP synergy score to combined doses of synchronous A485 and Venetoclax (peak ZIP 62.24) in 697*^WT^* cells. Viability measured by 72h MTS assay. **c,** Summary BODIPYC_11_ staining 697*^WT^* (black) and 697*^WT^* treated with A485 (pale blue) cells in response to Venetoclax. FL1:FL3 signal normalized to DMSO vehicle. Performed in triplicate, each dot represents a single sample, mean ± SD, two way ANOVA, ***, *P* <0.0006, ****, *P* <0.0001. **d,** Summary of basal (left) and maximal (middle) mitochondrial oxygen consumption rate (OCR) and spare respiratory capacity (right) measured using Seahorse (Agilent) Mitostress test in 697*^WT^* cells treated with A485 (pale blue) or DMSO vehicle (black). Each dot represents a single replicate acquired from two separate experiments. Unpaired *t* test ****, *P* <0.0001. **e,** Growth curve of 697*^WT^* cells treated with Venetoclax 100nM and/or A485 at two different doses (100nM and 300nM). Duplicate measurements, mean ± SD presented as log_2_ accumulative fold change. **f,** Annexin-V (APC) externalization assessed by flow cytometry in 697*^WT^*cells treated with Venetoclax 100nM and/or A485 at two different doses (100nM and 300nM). Duplicate measurements, each dot represents a single replicate, mean fluorescent intensity (MFI) ± SD normalized to DMSO control. One way ANOVA, ****, *P*<0.0001; ***, *P*=0.0004; **, *P*=0.027. **g,** Dose response curve of NALM6*^WT^* (solid line) and NALM6*^WT^* pre-treated with 3 days of A485 (hashed line) to Venetoclax in 72h MTS viability assays. Performed in triplicate, mean ± SD. **h,** Summary BODIPYC_11_ staining NALM6*^WT^* (red) and NALM6*^WT^* treated with A485 (brown) cells in response to Venetoclax. FL1:FL3 signal normalized to DMSO vehicle. Performed in triplicate, each dot represents a single sample, mean ± SD, two way ANOVA, ****, *P* <0.0001. **i,** Three-dimensional diffusion plot of ZIP synergy score to combined doses of synchronous A485 and Venetoclax. Viability measured by 72h MTS assays in the Venetoclax-sensitive cell lines SUPB15 (*BCR::ABL1*) (peak ZIP 13.86), KOPN8 (*MLL::ENL*) (peak ZIP 12.66) and RS4-11 (*MLL::AF4*) (peak ZIP 27.08).

We extended these findings to other human B-ALL cell lines. A485 sensitized the highly Venetoclax-resistant, *BAX*-mutated cell line NALM6 to Venetoclax (Fig. 5g). This was associated with increased lipid peroxidation, indicative of ferroptotic death (Fig. 5h and Extended Data Fig. 5h). In Venetoclax-sensitive B-ALL lines driven by high-risk genetic drivers, Venetoclax and A485 showed evidence of pharmacological synergy, indicating that this interaction is conserved across diverse genetic subtypes of B-ALL (Fig. 5i).

Collectively, these findings pharmacologically validate the findings of BCL2 sensitization identified in our *CREBBP* mutated cell lines and support a possible novel drug combination for clinical translation in B-ALL more broadly.

### Genetic or pharmacological inhibition of CREBBP sensitizes B-ALL to Venetoclax *in-vivo*

Finally, we sought to test whether Venetoclax could target *CREBBP*-mutated B-ALL *in-vivo*, where cell extrinsic factors and pharmacodynamic effects can result in reduced efficacy or highlight dose-limiting toxicities.

Luciferase-expressing 697*^WT^* and 697*^KI^*cell lines were engrafted into NOD-SCID-Gamma (NSG) mice (Extended Data Fig. 6a,b). Upon confirmation of engraftment by bioluminescent imaging (BLI), mice were treated with daily oral Venetoclax for up to 30d (Fig. 6a). As anticipated, Venetoclax exposure was associated with limited disease control in 697*^WT^* cells (Fig. 6b,c) resulting in a small, but significant, prolongation of post-transplant overall survival (OS), with all Venetoclax-treated 697*^WT^*recipient animals succumbing within 15d of treatment (median OS 19 vs. 22d, *p*=0.0011) (Fig. 6f). 697*^KI^* recipient mice treated with vehicle control succumbed to disease slightly later than 697*^WT^*, potentially reflecting the proliferation defect characterized *in-vitro* (median OS 25 vs. 19d, *p*=0.0004) (Fig. 6f); nevertheless, all vehicle-treated 697*^KI^* mice succumbed by 19d of treatment. Venetoclax-treated mice engrafted with 697*^KI^* cells demonstrated markedly improved disease control by BLI (Fig. 6d,e and Extended Data Fig. 6c,d). Only one Venetoclax-treated 697*^KI^* recipient succumbed to disease within the treatment window, with the remaining 5 recipients gaining up to 2 weeks of survival after cessation of Venetoclax exposure (median OS 47.5 vs. 25.5d, *p*=0.0006) (Fig. 6f).

**Figure 6:**
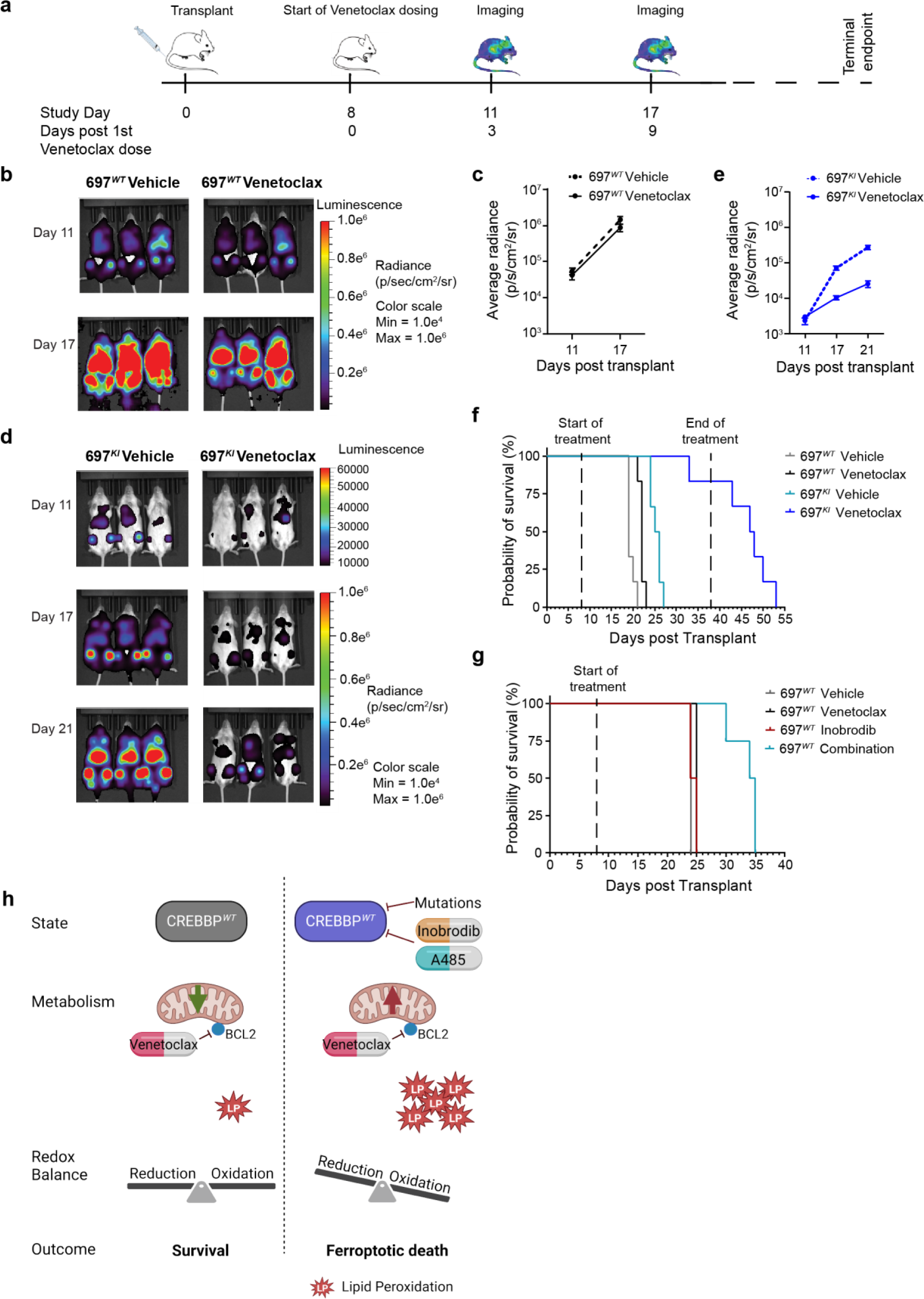
Genetic or pharmacological inhibition of CREBBP sensitizes B-ALL to Venetoclax *in-vivo*. **a,** Schema of 697*^WT^* vs. 697*^KI^ in-vivo* drug dosing protocol. NSG mice were intravenously injected with 697*^WT^*vs. 697*^KI^* luciferase-expressing cell lines. At 8 days post injection animals were administered Venetoclax (100mg/kg) or vehicle control by oral gavage. Mice were imaged by BLI as indicated. N=6 per group. **b,** Representative BLI images of mice engrafted with 697*^WT^* cells at days 11 and 17 of treatment. **c,** Average BLI radiance of mice engrafted with 697*^WT^* cells at days 11 and 17 of treatment (log p/s/cm^2^/sr). Mean ± SEM. **d,** Representative BLI images of mice engrafted with 697*^KI^* cells at days 11, 17 and 21 of treatment. **e,** Average BLI radiance of mice engrafted with 697*^KI^* cells at days 11, 17 and 21 of treatment (log p/s/cm^2^/sr). Mean ± SEM. **f,** Overall survival of NSG mice engrafted with 697*^WT^* vs. 697*^KI^* B-ALL cells treated with oral Venetoclax (100mg/kg) or vehicle control. 30 day maximal treatment window is indicated. N=6 per group. Significance calculated by Mantel Cox log-rank test. **g,** Overall survival of NSG mice engrafted with 697*^WT^* B-ALL cells treated with oral Inobrodib (20mg/kg) or a combination of Inobrodib (20mg/kg) plus Venetoclax (100mg/kg) (n=4 per group). Engraftment control groups were treated with Venetoclax only (n=2) or vehicle control (n=1). Treatment began at day 8 post-transplant. Significance calculated by Mantel Cox log-rank test. **h,** Summary of relationship between metabolism and redox balance in B-ALL in presence of normal CREBBP function (left) or *CREBBP* mutation or inhibition (right). For explanation, see also text in discussion.

Lastly, we tested whether co-administration of the orally-available, early-phase CREBBP bromodomain inhibitor Inobrodib could sensitize 697*^WT^* cells to Venetoclax *in-vivo* (Extended Data Fig. 6e). Once-daily oral dosing of single-agent Inobrodib did not confer survival benefit compared to recipients treated with either vehicle or single-agent Venetoclax, whereas recipient mice treated with both agents demonstrated a significant survival advantage (median OS combination 34.5 vs. Inobrodib only 24.5d, p=0.0084) (Fig. 6g).

Overall, these findings demonstrate that: i) oral Venetoclax can be highly efficacious in controlling *CREBBP*-mutated B-ALL in a preclinical *in-vivo* model; and ii) pharmacological inhibition of CREBBP is a tolerable and efficacious means of sensitizing B-ALL to Venetoclax *in-vivo*.

## Discussion

There is a pressing clinical need to develop novel treatments for patients with high-risk B-ALL, such as those harbouring mutations in *CREBBP*. Using a highly-curated panel of clinically-tractable small molecules in an isogenic human cell line model, we demonstrate that *CREBBP* mutation sensitizes cells to Dexamethasone, inhibitors of residual CREBBP/EP300 function and most potently BCL2 inhibition. *CREBBP* LOF increases metabolic rate and upon BCL2 inhibition results in ferroptotic cell death, which can be phenocopied by pharmacological CREBBP inhibition in genetically-diverse B-ALL cell lines, providing a readily translatable novel drug combination for B-ALL more broadly.

Our model shows a number of significant similarities to previously published work. The global transcriptional profile of our *CREBBP*-mutated cells strongly overlaps with those seen in murine *Crebbp*^-/-^ germinal-centre B-cells^25^. Other models of *CREBBP*-mutated B-ALL and B-cell lymphoma have implicated changes in cell cycle, metabolism, DNA damage response and apoptosis^3,10,19^. We also note that hypodiploid B-ALL, a genetic subtype highly associated with *CREBBP* mutations, has been linked with Venetoclax sensitivity^20^. Moreover, preliminary results from an early phase clinical trial of Venetoclax in relapsed pediatric malignancies (NCT03236857) have specifically highlighted responses in the small number of *CREBBP*-mutated B-ALL patients enrolled, providing supportive evidence that our results will be clinically translatable^28^.

Our *CREBBP*-mutated model did not show significant differential sensitivity to cytotoxic chemotherapy in the *ex vivo* setting. Our cell lines also showed a small but significant sensitization to Dexamethasone, suggesting that *CREBBP* LOF does not significantly alter responses to glucocorticoids^4,11,12^. Previous reports have shown variable association of *CREBBP* LOF with chemoresistance *in-vitro*^10,12,19^ and overall, our findings add to a growing body of evidence that CREBBP LOF does not in itself provoke significant cell-intrinsic chemoresistance.

A number of possible routes to chemoresistance remain. Recent reports have shown that LOF mutations affecting multiple epigenetic regulators across multiple malignancies can increase tolerance to environmental stress, a characteristic unlikely to read out in standard *in-vitro* culture conditions^29^. A prominent phenotype of our model was cell cycle retardation, demonstrable both transcriptionally and in *in-vitro* assays, and potentially resulted in the relatively delayed engraftment of 697*^KI^* cells in NSG mice. It is therefore possible that slower cell-cycle kinetics could result in *relative* resistance to predominantly S-phase-targeting cytotoxic chemotherapy, as has been shown in pre-clinical models of chemoresistance in B-ALL^30^. The mechanism underlying delayed cell cycle progression is likely to be multifactorial, but we note recent work implicating CREBBP/EP300’s role in maintaining enhancer acetylation marks during cell division^31^. Thus, a failure to efficiently reconstitute the enhancer landscape during mitosis could plausibly result in delayed cell-cycle progression and may also explain why genetic or pharmacologically-induced loss of CREBBP function exerts a more potent phenotype^4^ than complete loss of CREBBP protein, through steric competition with residual compensatory EP300 activity.

We show that BCL2 inhibition results in ferroptotic cell death in the context of *CREBBP* LOF. Unlike apoptosis, ferroptosis is not mediated by a defined biochemical pathway, rather is the output of a combination of underlying metabolic state, ROS scavenging capacity and lipid composition^32^. B-cell progenitors are exquisitely sensitive to redox balance^33^ and recent genetic perturbation data has highlighted an underlying propensity for ferroptotic cell death in B-ALL^34^. Reflecting its pleiotropic role in biology^35^, how *CREBBP* loss alters this balance is likely to be multifactorial, including by direct transcriptional changes, as well as post translational acetylation of proteins affecting metabolism and redox balance (Fig. 6h). Our functional experiments show significant increases in both glycolytic and mitochondrial metabolism upon CREBBP LOF. Thus, in contrast with studies in AML, we show an association between *enhanced* metabolic state and BCL2 dependence, likely relating to a B-cell-state specific susceptibility to oxidative stress^23,34^. Concordantly, we hypothesize that the deep repression of *AGPS* seen in 697*^KI^* cells is an adaptation to limit production of the unsaturated lipid substrates responsible for ferroptosis^36^.

Lastly, we demonstrate that small molecule inhibition of CREBBP can sensitize genetically-diverse B-ALL cell lines to Venetoclax, potentially widening the paradigm of our combination to multiple ALL genotypes, and the clinically-actionable combination of Venetoclax and Inobrodib was tolerable and highly efficacious in *in-vivo* models. We are aware of preliminary reports of this combination being trialled in related haematological malignancies^37^ and propose this as a rational drug combination for B-ALL.

## Methods

### Cell lines

B-ALL cell lines 697, REH and NALM6 cell lines were purchased from DSMZ. RS411, KOPN8 and SUPB15 were provided by Prof. Owen Williams (UCL, UK). 697, REH, KOPN8, RS4-11 were cultured at 37°C with 5% CO2 in IMDM (ThermoFisher, 12440053) supplemented with 10% v/v heat-inactivated fetal bovine serum (HI-FBS) (Sigma Aldrich, F7524), Penicillin-Streptomycin 100U/ml (Sigma Aldrich, P0781) and 1% v/v L-glutamine (Sigma Aldrich, G7513). NALM6 and SUPB15 were cultured in IMDM 20% serum. HEK-293T cells were maintained in DMEM (Sigma D0819), 10% FBS, L-glutamine, and penicillin/streptomycin. Cell lines were tested for mycoplasma by PCR (CSCI Core Facility) and STR genotyped using Promega Powerplex 16-HS kit (performed by Genetica LabCorp). Cells were routinely passaged to 1×10^6^/ml every 2-4 days.

### Generation of *CREBBP* Mutant Lines

697 and REH cells were lentivirally-transfected with Cas9, single cell cloned and tested for high Cas9 activity by GFP/BFP reporter assay. *CREBBP^R^*^1446^*^C^* gene-editing reagents were introduced by nucleofection (Nucleofector II, Kit R, Lonza) consisting of a gRNA/Cas9 RNP complex (IDT 1072532, 1081060: target sequence CATGGTAAACGGCTGTGCGG), two AltR ssDNA Ultramers silently mutating the PAM sequence with or without insertion of the R1446C mutation (50:50 mix: T*T*CACATACTCTAAATATCCAATAAGGATCTCATGGTAAACGGCTGTGCAGAGACAACGTGGCCGGAAGAA ATGAATACTATCCAGATAAGAAATGTA*C*A; T*T*CACATACTCTAAATATCCAATAAGGATCTCATGGTAAACGGCTGTGCGGAGACAACGTGGCCGGAAGAA ATGAATACTATCCAGATAAGAAATGTA*C*A), 3mM homology directed repair enhancer (IDT, 1081072) and IDT nucleofection enhancer oligonucleotide (IDT 1075915) according to manufacturer’s protocol. At 48h ATTO550^+ve^ single cells were FACS sorted into 96-well plates (BD FACSAria Fusion cell sorter) and, upon growth, colonies screened for successful editing by HpyCH4V amplicon digestion (F:GAGCACCTGGAAAGAGGAGC; R:CCCACAGGCGTGTGTACATT). Clones were validated by Sanger sequencing of the bulk screening amplicon and 10xTOPO-TA-cloned PCR products (ThermoFisher, K4575J10).

### MTS

All drugs were purchased from Selleckchem, except A1155463 (Cayman Chemicals), and reconstituted to 10mM in DMSO. 72-hour viability assays were performed using CellTiter 96® AQ_ueous_ One Solution Cell Proliferation Assay (Promega, G3580) following manufacturer recommendations and 490nM absorbance measured using a CLARIOstar plate reader (BMG Labtech). Drug synergy scores were calculated using SynergyFinder web tool (https://synergyfinder.fimm.fi)^38^. We defined a peak ZIP threshold of +10 for synergism.

### Mitochondrial Depolarization using JC1

Mitochondrial depolarization was assessed on cells exposed to Venetoclax for 24h using the MitoProbe^TM^ JC-1 assay kit (ThermoFisher, M34152) following manufacturer recommendations. Cells were analysed by flow cytometry (BD Fortessa) for 488nm-excitied 530/30 filter signal. Viable cells were gated by DAPI (BD, 564907) exclusion. During flow cytometry cells were routinely handled in DPBS (Gibco, 14190-094) 2% HI-FBS and 2mM EDTA (Invitrogen, 15575038).

### Annexin-V Staining by Flow Cytometry

Annexin-V externalization was assessed using anti-AnnexinV-APC kit from (eBioscience^TM^ 88-8007-74) following manufacturer protocol. Briefly, after incubation with drug/vehicle, cells were washed in DPBS, resuspended in Annexin binding buffer and incubated with 1/40 Annexin-V-APC dilution for 15min at room temperature (RT) in the dark. Cell pellet was resuspended in 7-AAD (BD, 559925) diluted 1/100 in 1x binding buffer and analyzed by flow cytometry.

### Western Blot

Protein lysates were obtained from 1×10^6^ cells after resuspending the pelleted cells in 50µl boiling lysis buffer (31mg/ml DTT, 10% SDS, 10% Glycerol, 1% Bromophenol blue and 12.5% 1MTris-HCL). Cell lysates were separated by SDS-PAGE electrophoresis (NuPAGE 3-8%TRIS acetate protein gels (Invitrogen) or 4-20% Mini-PROTEAN TGX protein gel (Bio-Rad)). Precision-plus protein dual color ladder (Bio-Rad, #1610374) or HiMark™ Pre-stained Protein Standard (Invitrogen, LC5699) were used as a size marker. Proteins were transferred to a methanol pre-activated membrane (Merck, IPFL00010) and then blocked with 5% BSA (ThermoFisher, BP1600) or powdered milk in TBS-0.05% Tween20 (TBST) before proceeding to antibody staining. Primary and secondary antibodies were diluted in 5% BSA or milk in TBST. Primary antibodies were incubated overnight at 4°C or 1h at RT; secondary antibodies were incubated for 1h at RT. Washes were performed with TBST. Membrane fluorescence was acquired using Odyssey imager (LI-COR).

Antibodies used: Caspase 9/PARP (Abcam, Ab136812), BCL2 (Abcam, ab32124), β-Tubulin (Sigma-Aldrich, T8328), CREBBP (Santa Cruz biotechnology, SC369) and Vinculin (Santa Cruz biotechnology, sc25336).

### Lentiviral Production

Lentiviral vectors were produced in HEK-293T cells by co-transfection of the previously sequenced vector constructs with psPAX and pMDG.2 packaging plasmids using Trans-IT LT-1 (Mirus, MIR2700). HEK293T cells were incubated overnight at 37°C and the next day medium was changed for IMDM 10% HI-FBS. Viral particles were harvested at 48 and 72h by centrifugation and 45μm filtration. 250µl of viral particles were used to transfect 1×10^6^ B-ALL cells by spinoculation (900g for 2h at 32°C) in the presence of 10μg/ml Polybrene (sc-134220). Subsequently, the cultures were diluted in fresh media and washed three times the following morning.

Transfection efficiencies were assessed by flow cytometric reporters and transfected cells selected using either 0.5μg/ml of Puromycin (Gibco) for three days or FACS cell sorting as appropriate to achieve >90% transfection.

### shRNA Knockdown of *BCL2*

Doxycycline-inducible reported shRNA knockdown constructs (pLT3GEPIR_GFP/Cherry_Stuffer) were provided by Dr Thomas Mercher (originally from Prof Johannes Zuber), *EcoRI/XhoI* double digested and the vector backbone purified by gel purification as previously described ^24^. *BCL2* targeting shRNAs were synthesized as Ultramers (IDT) based on the following recommended sequences (BCL2_1: TGCTGTTGACAGTGAGCGCCCGGGAGATAGTGATGAAGTATAGTGAAGCCACAGATGTATACTTCATCACTATCTCCCGGTTGCCTACTGCCTCGGA; BCL2: TGCTGTTGACAGTGAGCGCGAGGATCATGCTGTACTTAAATAGTGAAGCCACAGATGTATTTAAGTACAGCATGATCCTCTTGCCTACTGCCTCGGA; Renilla: TGCTGTTGACAGTGAGCGCAGGAATTATAATGCTTATCTATAGTGAAGCCACAGATGTATAGATAAGCATTATAATTCCTATGCCTACTGCCTCGGA) ^24^. Complementary oligonucleotides were resuspended to 500ng/µl, boiled and hybridized, ligated using T4 ligase (NEB) (5ng insert:50ng vector) and transformed into chemically-competent *E. coli*. Bacterial colonies were screened by PCR (F:CTCGACTAGGGATAACAGGG R:CAAAGAGATAGCAAGGTATTCAG) and successful plasmid clones miniprepped (Qiagen) and sequence verified by Sanger sequencing. Cells were selected with puromycin 0.5μg/ml. Cells were incubated with 500ng/ml of doxycycline and mCherry positive cells FACS sorted to confirm knock-down by qPCR and western blot.

Competitive co-culture assays were performed by mixing an equal number of *renilla* shRNA-GFP and one of two *BCL2* shRNA-mCherry transfected 697*^WT^* or 697*^KI^* cells in triplicate. *BCL2* knockdown was induced with doxycycline and aliquots of cell culture analyzed daily by flow cytometry. Results are presented as the ratio of 697*^KI^* Cherry:GFP events normalized to the mean ratio of 697*^WT^* Cherry:GFP events.

### Gene expression by Reverse Transcriptase Quantitative PCR (RT-qPCR)

Total RNA was extracted using the RNeasy kit (Qiagen) and 1μg was used to synthesize cDNA using TaqMan Reverse transcription (Applied Biosystems, N8080234) following manufacturer’s instructions. RT-qPCR was carried out with SYBR Green PCR master mix (Agilent, 600882) on cDNA (diluted 1:5 in water). The expression level of RNA was calculated using the standard curve method, normalized to the expression of *GAPDH*. Primer pairs: *BCL2*F:ATTGGGAAGTTTCAAATCAGC; *BCL2*R:TGCATTCTTGGACGAGGG; *GAPDH*F:CCACATCGCTCAGACACCAT; *GAPDH*R:CCAGGCGCCCAATACG.

### Cell Cycle Analysis by FUCCI Reporter

CSII-EF-MCS vectors encoding mCherry-hCdt1(30/120) and mVenus-hGeminin(1/110) were obtained from Atsushi Miyawaki, RIKEN BioResource Research Center ^39,40^. Lentiviral vectors were made and dual transfected into 697 isogenic cell lines as above. mCherry/mVenus dual-positive cells were enriched by FACS cell sorting (BD Influx) to generate high-expressing stable cell lines and analyzed by flow cytometry.

### Seahorse Metabolic Assays

Oxphos and glycolysis assays were performed using the Seahorse XFe96 analyser (Agilent) according to manufacturer’s instructions. Briefly, cells were analysed at 48-72h post passage and in drug concentrations/DMSO vehicle as stated in figure legends. For glycostress experiments, cells were harvested and counted in Seahorse XF base medium, pH7.4 supplemented with 2mM L-Glutamine (Sigma Aldrich, G7513). The final concentration of the injected drugs in the glycostress test is 10mM glucose (Sigma, G7021), 1µM Oligomycin A and 50mM of 2-deoxyglucose (2-DG) (Sigma, D3179). For mitostress tests, Seahorse XF base media was supplemented with 2mM L-Glutamine, 10mM Glucose and 1mM Sodium pyruvate (Sigma, S8636). The final concentration of the injected drugs was 1µM Oligomycin, 1µM FCCP supplemented with 1µM Sodium pyruvate and 1µM Antimycin. For experiments using Venetoclax or A485 pretreatment drug exposure continued in the Seahorse media. Cells were plated at a density of 70000 cells/well in XFp tissue culture plates previously coated with CellTak (Corning, 354240) and incubated for 45mins at 37°C in a CO2-free incubator prior to analysis.

### Lipid Peroxidation Assays

Lipid peroxidation was assessed using the lipid peroxidation sensor BODIPYC_11_-581/591 (ThermoFisher, D3861) according to manufacturer’s instructions. Briefly, cells were washed in DPBS and stained with 4µM BODIPY dye for 30 min at 37°C with gentle shaking in the dark. After washing with DPBS, cells were analyzed by flow cytometry. Replicate data is presented as the ratio of 488nm-excited 530/30 (“FL1”) to 610/20 (“FL3”) filtered signal.

### RNA Sequencing Library Preparation

697*^WT^* and 697*^KI^* treated with 20nM Venetoclax or DMSO vehicle for 24h (Fig 4) were harvested, washed and RNA was extracted from 4×10^6^ cells using the RNeasy kit (Qiagen) (performed in triplicate). Libraries were prepared using NEBNext Poly(A) mRNA magnetic Isolation module (NEB, E7490) starting with 1µg DNA-free RNA, as per manufacturer’s instructions. RNA fragmentation, double-strand cDNA synthesis, end repair, adapter ligation and PCR amplification (9 cycles) was performed using NEBNext Ultra II directional RNA library prep kit for Illumina protocol (NEB, E7760) according to manufacturer’s protocol. Library quality and molarity was measured by Qubit (ThermoFisher) and TapeStation (Agilent). Samples were 50-bp paired-end sequenced on the NovaSeq (Illumina) instrument (CRUK CI Genomics Core Facility).

### RNAseq analysis

The quality of the paired-end RNA-seq reads were assessed using FastQC (https://www.bioinformatics.babraham.ac.uk/projects/fastqc/). All RNA-seq libraries were found to be above the minimum quality thresholds across quality control metrics. Adapter sequences were trimmed using TrimGalore package (https://github.com/FelixKrueger/TrimGalore), and were next mapped against the human genome version 38 (Hg38) using STAR version=2.7.10a ^41^. Only uniquely-mapping high confidence reads (flags NH:i:1 and MAPQ=255) were retrieved and used for transcriptome quantification with featureCounts version 2.0.1 ^42^. Genes with a minimum of 10 reads or more in a minimum of two samples were considered for all downstream analysis. Differential expression analysis was based on either a single factor model (∼ genotype/treatment/sensitivity) or an interaction model when two factors were considered (∼ genotype * treatment), and the interaction term was used to determine the difference of the effect of treatment across genotypes. Differential expression analysis was conducted using gene level transcriptomic reads using the Bioconductor package DESeq2 ^43^. KEGG Pathway enrichment, Gene Set Enrichment Analyses (GSEA) and Gene Ontology (GO) Analyses were all performed using the R package ‘clusterProfiler’ version 4.4.4 ^44^. A minimum p-value (corrected for multiple testing) and FDR of 0.05 were used across all tests to establish statistical significance.

### *In-Vivo* dosing studies

For the *in-vivo* dosing experiments in Figure 7A-F, 0.25×10^6^ luciferase expressing cells (isogenic cell lines 697*^WT^* & 697*^KI^*) were injected by tail vein injection into sub-lethally (2Gy) irradiated 13-15 week-old female NSG (NOD.Cg-Prkdcscid Il2rgtm1Wjl/SzJ) mice. Mice received a daily dose of Venetoclax (100mg/kg)(LC Labs) or vehicle control (15% Kolliphor HS15 (Sigma 42966), 60% Phosal propylene glycol (MP, 151957) and 10% ethanol), by means of Oral Gavage (FTP-20-30, INSTECH) for thirty days or until the experimental endpoints were reached. Disease progression was tracked by IVIS bioluminescence imaging (PerkinElmer); in brief, D-luciferin (Perkin Elmer, 122799) was administered by intraperitoneal (IP) injection (10µl/g body weight of a 15mg/ml solution) followed by inhalation anesthesia (isoflurane) and IVIS bioluminescence imaging (∼10 minutes post-luciferin injection). Tumor burden was quantified using Living Image Software (v4.7.2, PerkinElmer).

For the *in-vivo* dosing experiments in Figure 7G, 0.25×10^6^ 697*^WT^* cells were injected by tail vein injection into sub-lethally (2Gy) irradiated 11-13 week-old old male NSG mice. Mice received a daily dose of Venetoclax (100mg/kg) and/or Inobrodib (20mg/kg) or vehicle control (as above), by means of oral gavage until the experimental endpoints were reached.

Mice were housed in a pathogen-free animal facility and were allowed unrestricted access to food and water. All experiments were conducted under a UK Home Office project (under the Animals (Scientific Procedures) Act 1986, Amendment Regulations (2012)) and following ethical review by the University of Cambridge Animal Welfare and Ethical Review Body.

### Statistics

Flow cytometric data was analyzed using FlowJo. Statistical analysis was performed in GraphPad Prism-v9 and Microsoft Excel. Seahorse data was processed on Wave software. Significance tests are stated in figure legends

### Data and Materials Availability

Sequencing data will be GEO deposited.

Further information and requests for resources and reagents should be directed to and will be fulfilled by the corresponding author.

## Acknowledgments

SER is supported by a Clinician Scientist Fellowship from Cancer Research UK (C67279/A27957) and Leukaemia UK John Goldman Fellowship (2022/JGF/004). We would like to thank funders of the Huntly laboratory including Cancer Research UK (C18680/A25508, C355/A26819 and DRCRPG-Nov22/100014), the European Research Council (647685), MRC (MR-R009708-1 and MR-X008371) and the Kay Kendall Leukaemia Fund (KKL1243 and KKL1440). This research was supported by the NIHR Cambridge Biomedical Research Centre (BRC-1215-20014) and the Cancer Research UK Cambridge Centre (Cancer Research UK Major Centre Award C9685/A25117), and was funded in part by the Wellcome Trust, who supported the Wellcome – MRC Cambridge Stem Cell Institute (203151/Z/16/Z) and Cambridge Institute for Medical Research (100140/Z/12/Z). The views expressed are those of the authors and not necessarily those of the NIHR or the Department of Health and Social Care. For the purpose of Open Access, the author has applied for a CC BY public copyright licence to any Author Accepted Manuscript version arising from this submission. We acknowledge support from the Cancer Research UK Cambridge Centre Genomics Facility, the Cambridge Institute for Medical Research Flow Cytometry Facility and the NIHR Cambridge BRC Cell Phenotyping Hub. We thank Thomas Mercher and Johannes Zuber for use of the pLT3GEPIR inducible shRNA system. We thank Atsushi Miyawaki at the RIKEN BioResource Research Center for the use of the FUCCI cell cycle reporter system.

## References

1 Hunger, S. P. & Mullighan, C. G. Acute Lymphoblastic Leukemia in Children. N Engl J Med 373, 1541–1552 (2015). 10.1056/NEJMra1400972

2 Pasqualucci, L. et al. Inactivating mutations of acetyltransferase genes in B-cell lymphoma. Nature 471, 189–195 (2011). 10.1038/nature09730

3 Horton, S. J. et al. Early loss of Crebbp confers malignant stem cell properties on lymphoid progenitors. Nat Cell Biol 19, 1093–1104 (2017). 10.1038/ncb3597

4 Mullighan, C. G. et al. CREBBP mutations in relapsed acute lymphoblastic leukaemia. Nature 471, 235–239 (2011). 10.1038/nature09727

5 Brady, S. W. et al. The genomic landscape of pediatric acute lymphoblastic leukemia. Nat Genet 54, 1376–1389 (2022). 10.1038/s41588-022-01159-z

6 Holmfeldt, L. et al. The genomic landscape of hypodiploid acute lymphoblastic leukemia. Nat Genet 45, 242–252 (2013). 10.1038/ng.2532

7 Inthal, A. et al. CREBBP HAT domain mutations prevail in relapse cases of high hyperdiploid childhood acute lymphoblastic leukemia. Leukemia 26, 1797–1803 (2012). 10.1038/leu.2012.60

8 Li, B. et al. Therapy-induced mutations drive the genomic landscape of relapsed acute lymphoblastic leukemia. Blood 135, 41–55 (2020). 10.1182/blood.2019002220

9 Waanders, E. et al. Mutational landscape and patterns of clonal evolution in relapsed pediatric acute lymphoblastic leukemia. Blood Cancer Discov 1, 96–111 (2020). 10.1158/0008-5472.BCD-19-0041

10 Gao, C. et al. Downregulating CREBBP inhibits proliferation and cell cycle progression and induces daunorubicin resistance in leukemia cells. Mol Med Rep 22, 2905–2915 (2020). 10.3892/mmr.2020.11347

11 Chougule, R. A., Shah, K., Moharram, S. A., Vallon-Christersson, J. & Kazi, J. U. Glucocorticoid-resistant B cell acute lymphoblastic leukemia displays receptor tyrosine kinase activation. NPJ Genom Med 4, 7 (2019). 10.1038/s41525-019-0082-y

12 Dixon, Z. A. et al. CREBBP knockdown enhances RAS/RAF/MEK/ERK signaling in Ras pathway mutated acute lymphoblastic leukemia but does not modulate chemotherapeutic response. Haematologica 102, 736–745 (2017). 10.3324/haematol.2016.145177

13 Lamble, A. J. et al. CREBBP alterations are associated with a poor prognosis in de novo AML. Blood 141, 2156–2159 (2023). 10.1182/blood.2022017545

14 Liu, Y. et al. An easy-to-use nomogram predicting overall survival of adult acute lymphoblastic leukemia. Front Oncol 12, 977119 (2022). 10.3389/fonc.2022.977119

15 Qu, X. et al. Genomic alterations important for the prognosis in patients with follicular lymphoma treated in SWOG study S0016. Blood 133, 81–93 (2019). 10.1182/blood-2018-07-865428

16 Malinowska-Ozdowy, K. et al. KRAS and CREBBP mutations: a relapse-linked malicious liaison in childhood high hyperdiploid acute lymphoblastic leukemia. Leukemia 29, 1656–1667 (2015). 10.1038/leu.2015.107

17 Meyer, S. N. et al. Unique and Shared Epigenetic Programs of the CREBBP and EP300 Acetyltransferases in Germinal Center B Cells Reveal Targetable Dependencies in Lymphoma. Immunity 51, 535–547 e539 (2019). 10.1016/j.immuni.2019.08.006

18 Lord, C. J., Quinn, N. & Ryan, C. J. Integrative analysis of large-scale loss-of-function screens identifies robust cancer-associated genetic interactions. Elife 9 (2020). 10.7554/eLife.58925

19 Dutto, I., Scalera, C. & Prosperi, E. CREBBP and p300 lysine acetyl transferases in the DNA damage response. Cell Mol Life Sci 75, 1325–1338 (2018). 10.1007/s00018-017-2717-4

20 Diaz-Flores, E. et al. Bcl-2 Is a Therapeutic Target for Hypodiploid B-Lineage Acute Lymphoblastic Leukemia. Cancer Res 79, 2339–2351 (2019). 10.1158/0008-5472.CAN-18-0236

21 Roca-Portoles, A. et al. Venetoclax causes metabolic reprogramming independent of BCL-2 inhibition. Cell Death Dis 11, 616 (2020). 10.1038/s41419-020-02867-2

22 Pollyea, D. A. et al. Venetoclax with azacitidine disrupts energy metabolism and targets leukemia stem cells in patients with acute myeloid leukemia. Nat Med 24, 1859–1866 (2018). 10.1038/s41591-018-0233-1

23 Lagadinou, E. D. et al. BCL-2 inhibition targets oxidative phosphorylation and selectively eradicates quiescent human leukemia stem cells. Cell Stem Cell 12, 329–341 (2013). 10.1016/j.stem.2012.12.013

24 Fellmann, C. et al. An optimized microRNA backbone for effective single-copy RNAi. Cell Rep 5, 1704–1713 (2013). 10.1016/j.celrep.2013.11.020

25 Zhang, J. et al. The CREBBP Acetyltransferase Is a Haploinsufficient Tumor Suppressor in B-cell Lymphoma. Cancer Discov 7, 322–337 (2017). 10.1158/2159-8290.CD-16-1417

26 Dixon, S. J. et al. Ferroptosis: an iron-dependent form of nonapoptotic cell death. Cell 149, 1060–1072 (2012). 10.1016/j.cell.2012.03.042

27 Zhao, J. et al. Human hematopoietic stem cell vulnerability to ferroptosis. Cell 186, 732–747 e716 (2023). 10.1016/j.cell.2023.01.020

28 Place, A. E. et al. Pediatric Patients with Relapsed/Refractory Acute Lymphoblastic Leukemia Harboring Heterogeneous Genomic Profiles Respond to Venetoclax in Combination with Chemotherapy. Blood 136 (2020). 10.1182/blood-2020-137376

29 Loukas, I. et al. Selective advantage of epigenetically disrupted cancer cells via phenotypic inertia. Cancer Cell 41, 70–87 e14 (2023). 10.1016/j.ccell.2022.10.002

30 Turati, V. A. et al. Chemotherapy induces canalization of cell state in childhood B-cell precursor acute lymphoblastic leukemia. Nat Cancer 2, 835–852 (2021). 10.1038/s43018-021-00219-3

31 Kikuchi, M. et al. Epigenetic mechanisms to propagate histone acetylation by p300/CBP. Nat Commun 14, 4103 (2023). 10.1038/s41467-023-39735-4

32 Dixon, S. J. & Pratt, D. A. Ferroptosis: A flexible constellation of related biochemical mechanisms. Mol Cell 83, 1030–1042 (2023). 10.1016/j.molcel.2023.03.005

33 Borzillo, G. V., Ashmun, R. A. & Sherr, C. J. Macrophage lineage switching of murine early pre-B lymphoid cells expressing transduced fms genes. Mol Cell Biol 10, 2703–2714 (1990). 10.1128/mcb.10.6.2703-2714.1990

34 Lalonde, M. E. et al. Genome-wide CRISPR screens identify ferroptosis as a novel therapeutic vulnerability in acute lymphoblastic leukemia. Haematologica 108, 382–393 (2023). 10.3324/haematol.2022.280786

35 Blobel, G. A. CREB-binding protein and p300: molecular integrators of hematopoietic transcription. Blood 95, 745–755 (2000).

36 Zou, Y. et al. Plasticity of ether lipids promotes ferroptosis susceptibility and evasion. Nature 585, 603–608 (2020). 10.1038/s41586-020-2732-8

37 Brooks, N. et al. CCS1477, a Novel p300/CBP Bromodomain Inhibitor, Enhances Efficacy of Azacitidine and Venetoclax in Pre-Clinical Models of Acute Myeloid Leukaemia and Lymphoma. Blood 138 (2021). 10.1182/blood-2021-148295

38 Zheng, S. et al. SynergyFinder Plus: Toward Better Interpretation and Annotation of Drug Combination Screening Datasets. Genomics Proteomics Bioinformatics 20, 587–596 (2022). 10.1016/j.gpb.2022.01.004

39 Sakaue-Sawano, A., Kobayashi, T., Ohtawa, K. & Miyawaki, A. Drug-induced cell cycle modulation leading to cell-cycle arrest, nuclear mis-segregation, or endoreplication. BMC Cell Biol 12, 2 (2011). 10.1186/1471-2121-12-2

40 Sakaue-Sawano, A. et al. Tracing the silhouette of individual cells in S/G2/M phases with fluorescence. Chem Biol 15, 1243–1248 (2008). 10.1016/j.chembiol.2008.10.015

41 Dobin, A. et al. STAR: ultrafast universal RNA-seq aligner. Bioinformatics 29, 15–21 (2013). 10.1093/bioinformatics/bts635

42 Liao, Y., Smyth, G. K. & Shi, W. featureCounts: an efficient general purpose program for assigning sequence reads to genomic features. Bioinformatics 30, 923–930 (2014). 10.1093/bioinformatics/btt656

43 Love, M. I., Huber, W. & Anders, S. Moderated estimation of fold change and dispersion for RNA-seq data with DESeq2. Genome Biol 15, 550 (2014). 10.1186/s13059-014-0550-8

44 Yu, G., Wang, L. G., Han, Y. & He, Q. Y. clusterProfiler: an R package for comparing biological themes among gene clusters. OMICS 16, 284–287 (2012). 10.1089/omi.2011.0118

