## Extended Data Table 1 and Extended Data Figures 1 to 6 for "Genetic or pharmacological inactivation of CREBBP sensitizes B-cell Acute Lymphoblastic Leukemia to Ferroptotic Cell Death upon BCL2 Inhibition"

| Drug | Target | Class | 697 WT | | 697 KI | | | 697 KO | | |
|---|---|---|---|---|---|---|---|---|---|---|
| | | | IC50 (M) | R squared | IC50 (M) | R squared | Mut:WT | IC50 (M) | R squared | Mut:WT |
| **Venetoclax** | **BCL2** | **Apoptosis** | 2.46E-06 | 0.8036 | 1.47E-08 | 0.9634 | 167.53 | 6.57E-08 | 0.9591 | 37.41 |
| **AZD5991** | **MCL1** | **Apoptosis** | 1.59E-06 | 0.9592 | 1.27E-06 | 0.9559 | 1.25 | 1.14E-06 | 0.932 | 1.39 |
| **Alisertib** | **AURKA** | **Cell Cycle** | 9.91E-08 | 0.847 | 1.05E-07 | 0.9324 | 0.95 | 8.38E-08 | 0.894 | 1.18 |
| **Palbociclib** | **CDK6** | **Cell Cycle** | 1.12E-07 | 0.9376 | 2.15E-07 | 0.9357 | 0.52 | 1.75E-07 | 0.9336 | 0.64 |
| **Cytarabine** | **Cytotoxic** | **Chemotherapy** | 1.64E-08 | 0.962 | 1.46E-08 | 0.9385 | 1.12 | 9.19E-09 | 0.9804 | 1.79 |
| **Daunorubicin** | **Cytotoxic** | **Chemotherapy** | 1.11E-08 | 0.9637 | 5.93E-09 | 0.9589 | 1.87 | 6.76E-09 | 0.9867 | 1.64 |
| **Vincristine** | **Cytotoxic** | **Chemotherapy** | 5.89E-10 | 0.9341 | 8.68E-10 | 0.8976 | 0.68 | 5.92E-10 | 0.9127 | 1.00 |
| **Dexamethasone** | **Glucocorticoid** | **Chemotherapy** | 2.35E-08 | 0.9154 | 5.39E-09 | 0.9402 | 4.37 | 8.66E-09 | 0.974 | 2.72 |
| **KU-55933** | **ATM** | **DNA Damage** | 2.08E-05 | 0.8681 | 2.98E-05 | 0.7868 | 0.70 | 6.78E-05 | 0.7902 | 0.31 |
| **Ceralasertib** | **ATR** | **DNA Damage** | 7.23E-07 | 0.9814 | 8.04E-07 | 0.9379 | 0.90 | 6.99E-07 | 0.979 | 1.03 |
| **MK8776** | **CHK1** | **DNA Damage** | 1.27E-06 | 0.9599 | 1.04E-06 | 0.9739 | 1.22 | 8.49E-07 | 0.9624 | 1.50 |
| **AZD7762** | **CHK1/2** | **DNA Damage** | 2.61E-07 | 0.6081 | 4.24E-07 | 0.7708 | 0.62 | 5.50E-07 | 0.8127 | 0.48 |
| **Olaparib** | **PARP** | **DNA Damage** | 4.10E-06 | 0.9767 | 2.33E-06 | 0.8915 | 1.76 | 3.23E-06 | 0.9238 | 1.27 |
| **ICG-001** | **CBP/Catenin** | **Epigenetic** | 2.94E-06 | 0.6648 | 6.54E-06 | 0.7572 | 0.45 | 2.95E-06 | 0.7918 | 1.00 |
| **Inobrodib** | **CREBBP/EP300 bromodomain** | **Epigenetic** | 1.26E-06 | 0.9789 | | | | 2.91E-07 | 0.9488 | 4.33 |
| **Inobrodib (fine dilutions *KI* only)** | **CREBBP/EP300 bromodomain** | **Epigenetic** | 4.61E-07 | 0.9832 | 7.00E-08 | 0.9619 | 6.58 | | | |
| **A485** | **CREBBP/EP300 HAT** | **Epigenetic** | 2.01E-07 | 0.8079 | | | | 4.56E-08 | 0.9731 | 4.42 |
| **A485 (fine dilutions *KI* only)** | **CREBBP/EP300 HAT** | **Epigenetic** | 2.77E-07 | 0.9164 | 6.94E-08 | 0.9558 | 3.99 | | | |
| **Tazemetostat** | **EZH2** | **Epigenetic** | No Reponse | NA | No Response | NA | NA | No Reponse | NA | NA |
| **Panobinostat** | **HDAC** | **Epigenetic** | 2.97E-09 | 0.9821 | 3.59E-09 | 0.9732 | 0.83 | 7.00E-09 | 0.9551 | 0.43 |
| **BRD3308** | **HDAC3** | **Epigenetic** | 8.75E-06 | 0.8207 | 7.14E-05 | 0.6267 | 0.12 | 8.97E-05 | 0.4664 | 0.10 |
| **IACS-010759** | **Mitochondrial Complex 1** | **Metabolic** | 2.91E-08 | 0.6432 | 1.53E-07 | 0.4615 | 0.19 | 4.64E-08 | 0.5464 | 0.63 |
| **Bortezomib** | **Proteosome** | **Proteosome** | 5.75E-07 | 0.991 | 5.68E-07 | 0.9878 | 1.01 | 5.41E-07 | 0.8512 | 1.06 |
| **Dasatinib** | **ABL1** | **Signaling** | 4.60E-07 | 0.9615 | 4.94E-07 | 0.9037 | 0.93 | 7.10E-07 | 0.9769 | 0.65 |
| **Ibrutinib** | **BTK** | **Signaling** | 1.30E-05 | 0.94 | 2.30E-05 | 0.9377 | 0.56 | 1.84E-05 | 0.9547 | 0.71 |
| **666-15** | **CREB** | **Signaling** | 1.76E-06 | 0.8966 | 1.28E-06 | 0.8698 | 1.38 | 9.02E-07 | 0.9163 | 1.95 |
| **Ruxolitinib** | **JAK-STAT** | **Signaling** | 1.10E-05 | 0.8882 | 8.51E-06 | 0.9284 | 1.29 | 2.28E-05 | 0.9261 | 0.48 |
| **SP600125** | **JNK** | **Signaling** | 5.83E-06 | 0.8064 | 1.66E-05 | 0.9055 | 0.35 | 9.80E-06 | 0.9243 | 0.59 |
| **BLZ945** | **MCSFR** | **Signaling** | No Reponse | NA | 3.87E-05 | 0.6412 | NA | 1.41E-04 | 0.5931 | NA |
| **Selumetinib** | **MEK** | **Signaling** | 2.65E-02 | 0.6962 | No response | NA | NA | 2.08E-03 | 0.7175 | 12.78 |
| **Idelalisib** | **PI3KD** | **Signaling** | 4.33E-05 | 0.5694 | 6.21E-05 | 0.8847 | 0.70 | 4.91E-05 | 0.6993 | 0.88 |
| **Dabrafenib** | **RAF** | **Signaling** | 5.65E-07 | 0.9359 | 4.61E-07 | 0.9219 | 1.23 | 3.48E-07 | 0.9536 | 1.62 |
| **RO5126766** | **RAF/MEK** | **Signaling** | 1.43E-05 | 0.8234 | 1.97E-05 | 0.7241 | 0.72 | 6.92E-06 | 0.8533 | 2.06 |
| **Lanraplenib** | **SYK** | **Signaling** | 1.39E-05 | 0.9385 | 1.86E-05 | 0.8192 | 0.75 | 1.97E-05 | 0.8872 | 0.71 |

**Extended Data Table 1:** Cell viability data from small molecule screen in $697^{WT}$, $697^{KI}$ and $697^{KO}$ cell lines (related to figure 1).

Extended Data Figure 1: *CREBBP*-mutated B-ALL shows increased sensitivity to Venetoclax

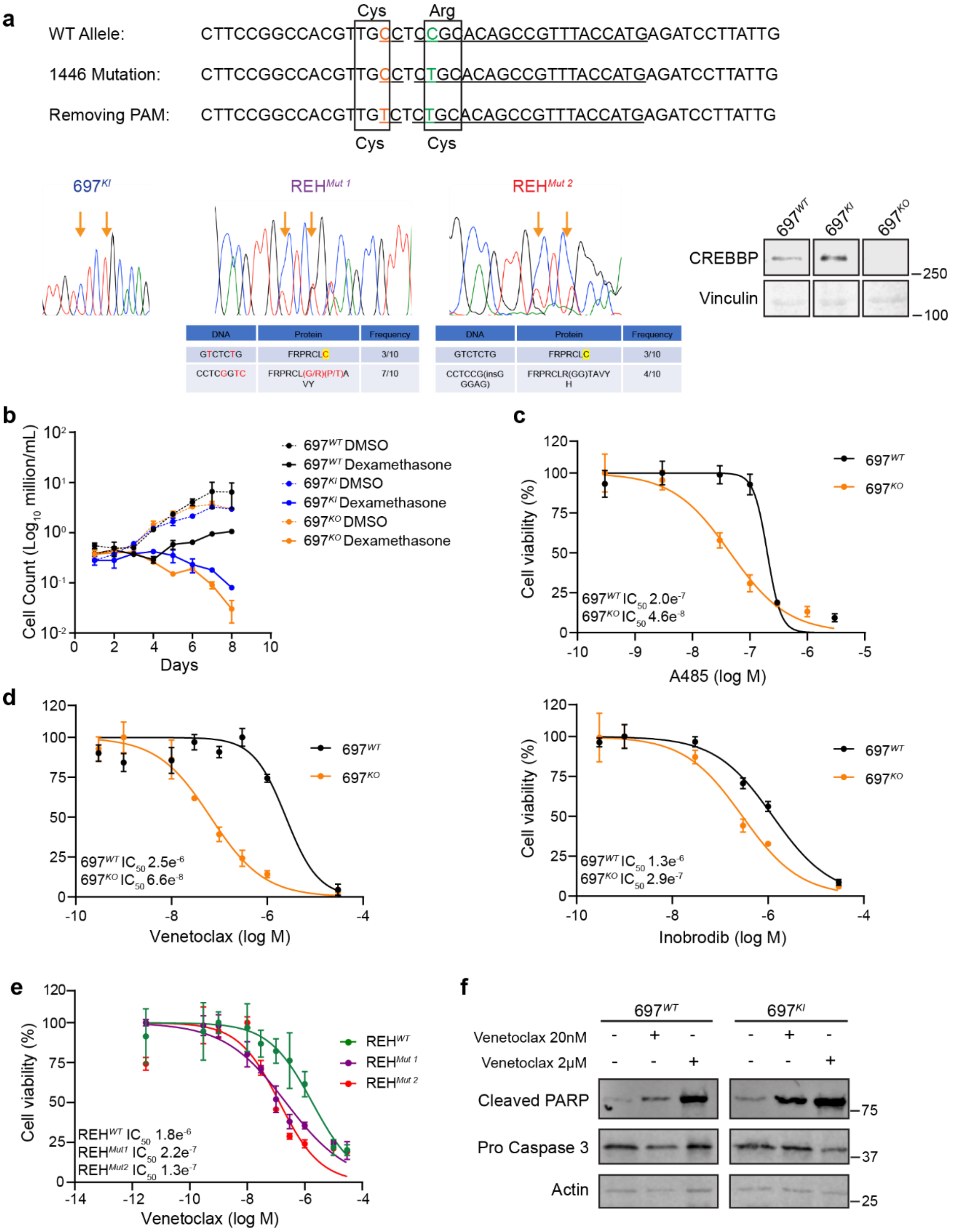


**Extended Data Figure 1: *CREBBP*-mutated B-ALL shows increased sensitivity to Venetoclax.**
**a,** Summary of genome editing strategy. Top: Two single base substitutions were introduced by CRISPR directed homologous recombination to: i) generate the R1446C mutation; and ii) remove the protospacer adjacent motif (PAM) to prevent further Cas9 binding and repeat cutting of successful edits. Bottom left: Results of amplicon sequencing showing a homozygous edit in $697^{KI}$ cells and a compound heterozygous edit in two REH mutant clones. The alternative sequences for the remaining two alleles were sequenced on TOPO-TA cloned amplicon fragments (sequences, amino acid substitution and TOPO-TA clonal frequency are shown in table below). Bottom right: Western blot of

CREBBP vs Vinculin protein in 697 edited clones, confirming loss of protein in $697^{KO}$ cells. Marker sizes in kDa shown.

**b,** Growth curves of $697^{WT}$ (black), $697^{KI}$ (blue) and $697^{KO}$ (yellow) grown in the presence of DMSO vehicle (hashed lines) or 10nM Dexamethasone (solid lines).

**c,** Dose response curves of two CREBBP/EP300 inhibitors A485 (top) and Inobrodib (bottom) showing enhanced sensitivity of $697^{KO}$ (yellow) compared to $697^{WT}$ (black) in 72h MTS viability assay. Performed in triplicate, mean ± SD.

**d,** Dose response curve of $697^{WT}$ (black) and $697^{KO}$ (yellow) lines to Venetoclax in 72h MTS viability assay. Performed in triplicate, mean ± SD.

**e,** Dose response curve of $REH^{WT}$ (green) and two isogenic *CREBBP*-mutant clones (purple and red) to Venetoclax in 72h MTS viability assays. Performed in triplicate, mean ± SD.

**f,** Western blot for cleaved PARP (top) and cleavage of pro-caspase 3 (middle) in $697^{WT}$ (left) and $697^{KI}$ (right) cell lines. Cells were incubated with DMSO vehicle or Venetoclax at either 20nM or 2000nM concentrations. Actin is shown as a loading control (bottom).

Extended Data Figure 2: Venetoclax exerts its effect on *CREBBP*-mutated B-ALL by on-target inhibition of BCL2

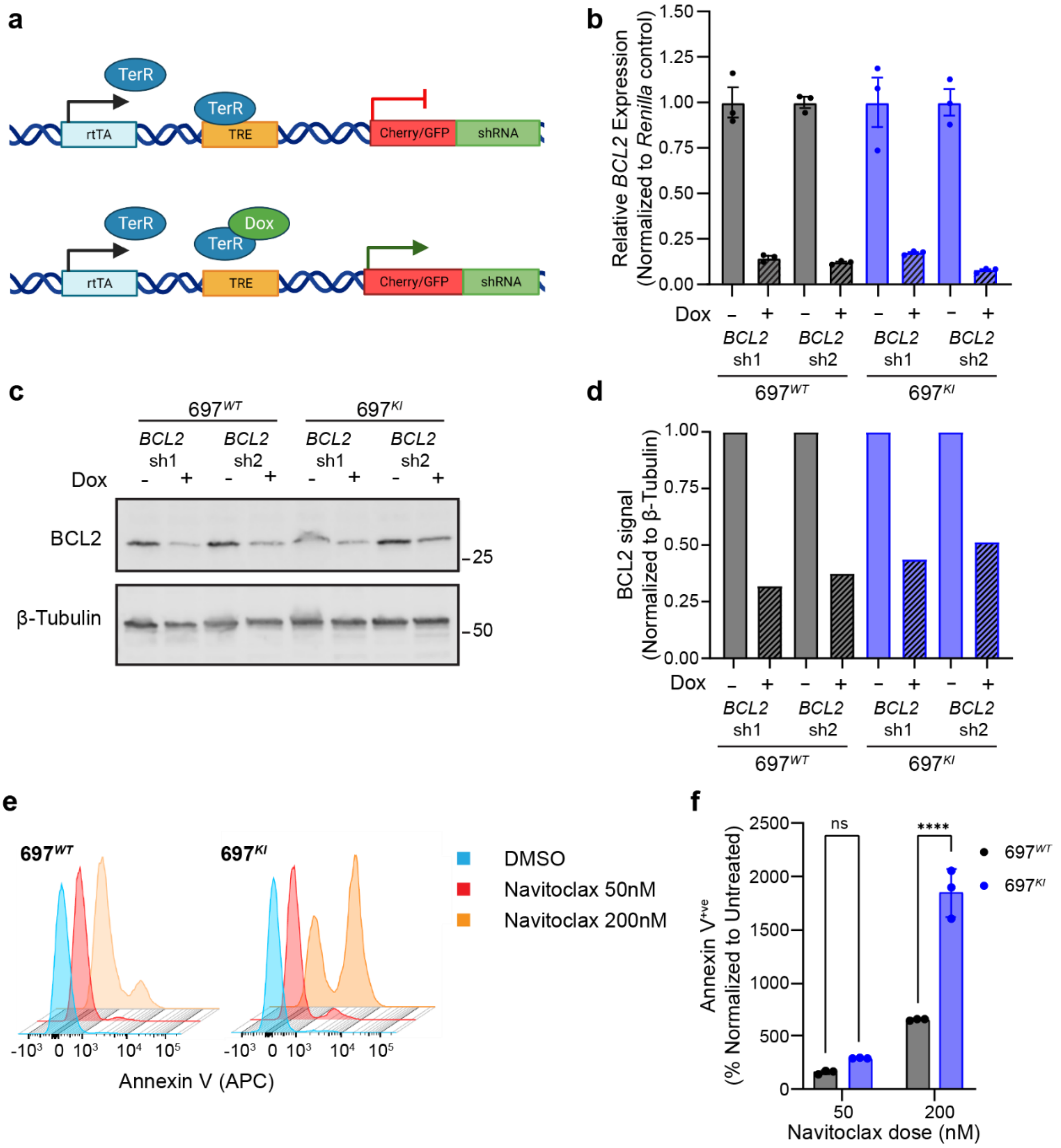


**Extended Data Figure 2: Venetoclax exerts its effect on *CREBBP*-mutated B-ALL on-target inhibition of BCL2.**

**a,** Schematic of doxycycline-inducible shRNA KD system linked to fluorescent reporter proteins [24]. Created with BioRender.com.

**b,** Doxycycline-induced KD of two different *BCL2*-targeting shRNAs measured by RT-qPCR. Triplicate measurements, internally normalised to *GAPDH* and presented as a ratio to *Renilla* control. Day 3 post induction. Mean ± SEM.

**c,** Western blot of BCL2 KD by two different doxycycline-inducible shRNAs in $697^{WT}$ (left) and $697^{KI}$ (right) cells. Day 3 post induction. Beta-Tubulin is presented as a loading control.

**d,** Secondary antibody fluorescence intensity from Fig. S2C normalized to Beta-Tubulin loading control.

**e,** Representative flow cytometry histograms of Annexin-V (APC) externalization in response to escalating doses of Navitoclax in $697^{WT}$ (left) and $697^{KI}$ (right).

**f,** Summary of replicate experiments in Fig. S2E measuring Annexin-$V^{+ve}$ cells normalized to DMSO-treated vehicle control cells. Performed in triplicate, mean ± SD, 2 way ANOVA $^{****}$, $P$ <0.0001.

Extended Data Figure 3: *CREBBP*-mutated B-ALL shows significant cell cycle and metabolic dysregulation

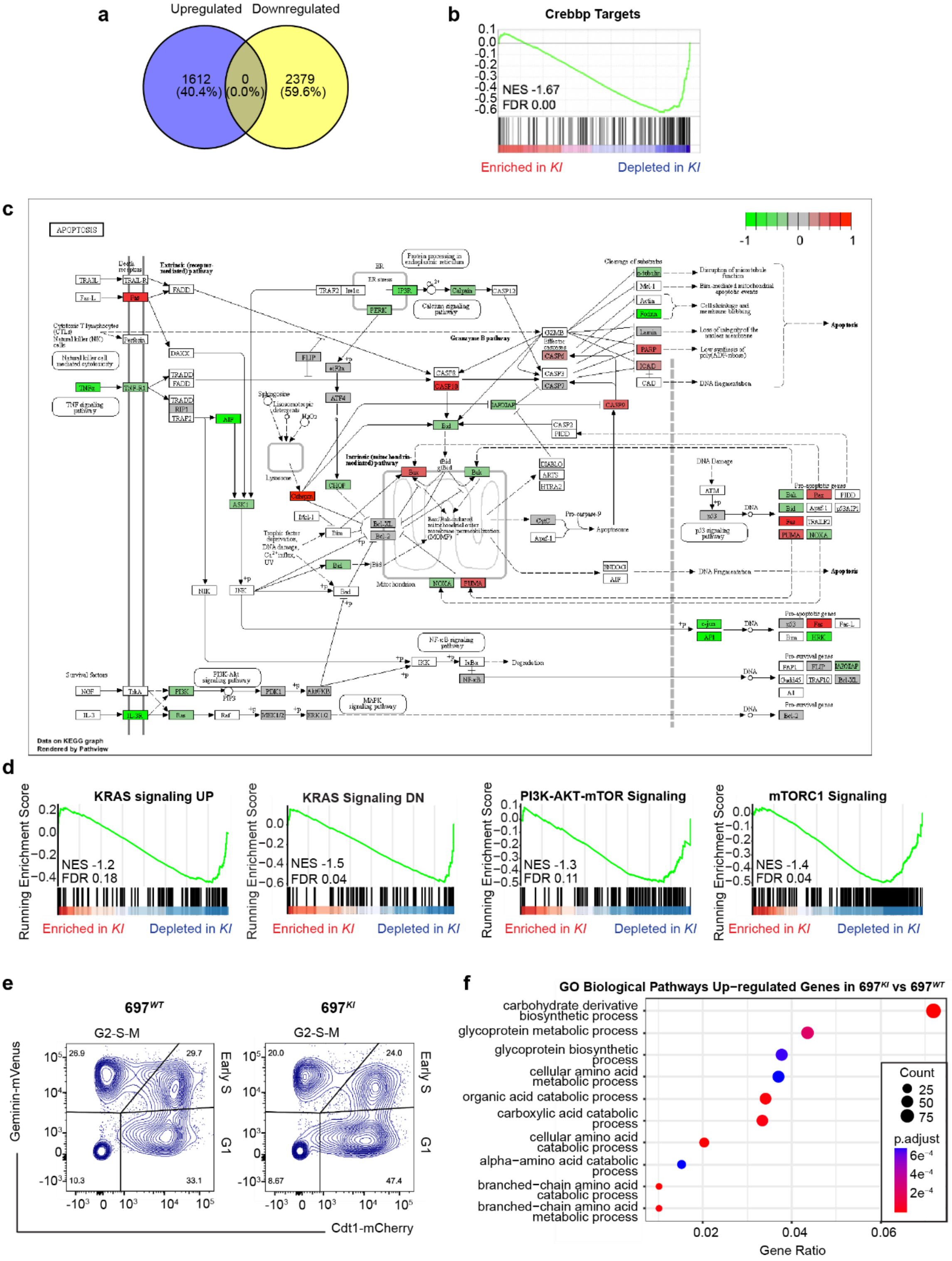


**Extended Data Figure 3: *CREBBP*-mutated B-ALL shows significant cell cycle and metabolic dysregulation.**

**a,** Number of up- and down-regulated DEGs (P and FDR <0.05) by RNAseq comparing $697^{KI}$ with $697^{WT}$ DMSO vehicle-treated cells.

**b,** GSEA of ranked RNAseq expression of $697^{KI}$ versus $697^{WT}$ DMSO vehicle-treated cells for known Crebbp target genes in *Crebbp*$^{KO}$ mouse germinal centre lymphocytes [25].

**c,** KEGG pathway showing differential expression of apoptotic regulators from RNAseq comparing $697^{KI}$ versus $697^{WT}$ DMSO vehicle-treated cells. Red genes upregulated, green downregulated.

**d,** GSEA of ranked RNAseq expression of $697^{KI}$ versus $697^{WT}$ DMSO vehicle-treated cells.

**e,** Representative plot of cell cycle stage by FUCCI reporter system in $697^{WT}$ (left) vs. $697^{KI}$ (right). Percentage viable single cells.

**f,** Significant up-regulated pathways identified by Gene Ontology (GO) database analysis of RNAseq comparing $697^{KI}$ versus $697^{WT}$ DMSO vehicle-treated cells.

Extended Data Figure 4: Venetoclax induces ferroptotic cell death in *CREBBP*-mutated B-ALL

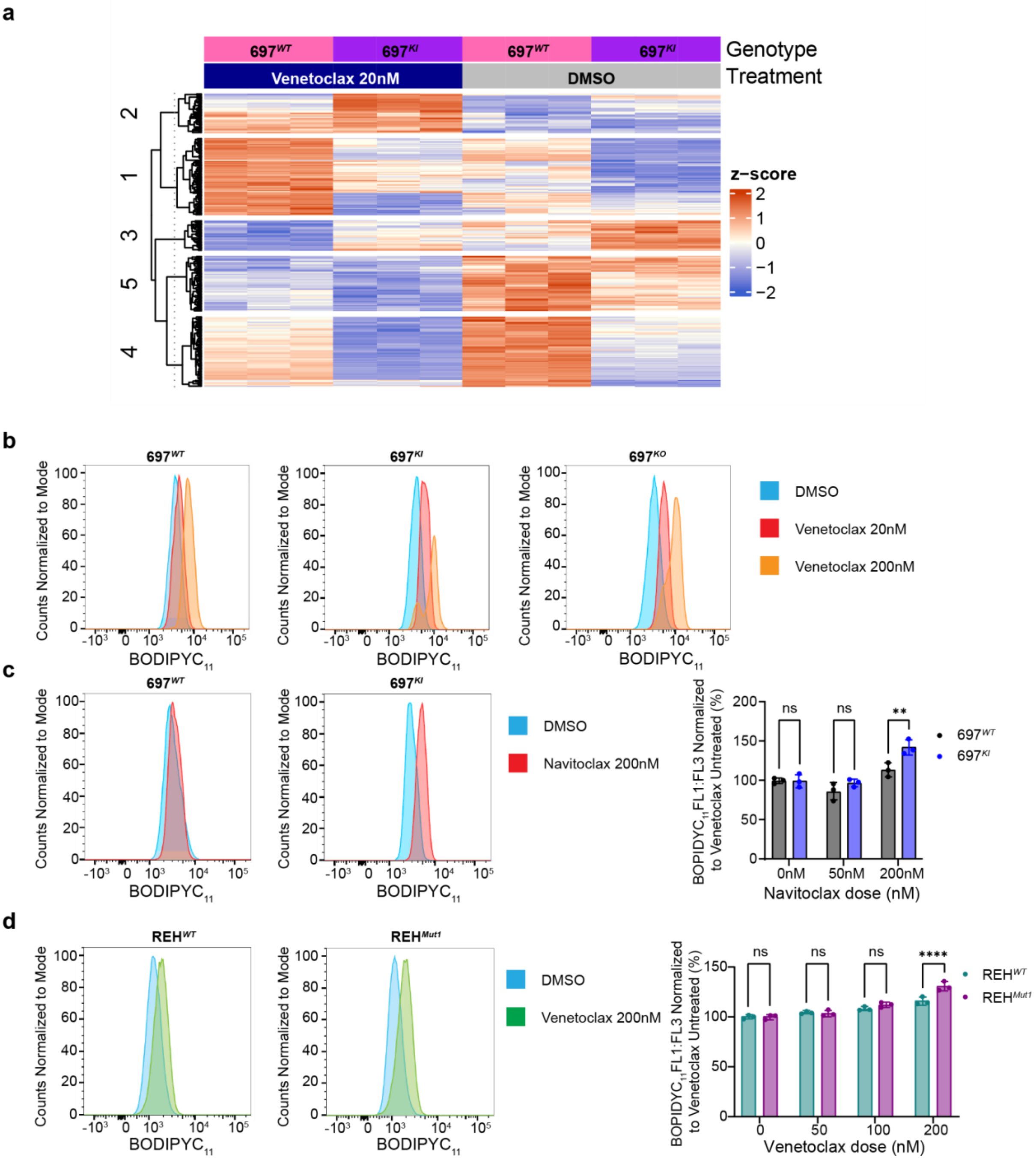


**Extended Data Figure 4: Venetoclax induces ferroptotic cell death in *CREBBP*-mutated B-ALL.**

**a,** Four-way interaction model identifies 1487 genes (FDR 0.05) differentially expressed specifically in Venetoclax-treated $697^{KI}$ cells.

**b,** Representative histogram of flow cytometric $BODIPYC_{11}$ staining (488nm 530/30) of $697^{WT}$ (left), $697^{KI}$ (middle) and $697^{KO}$ (right) cells in response to increasing doses of Venetoclax (DMSO: blue; Venetoclax 20nM: red; Venetoclax 200nM: orange).

**c,** Left panel: representative histogram of flow cytometric $BODIPYC_{11}$ staining (488nm 530/30) of $697^{WT}$ (left) and $697^{KI}$ (right) cells in response to 200nM Navitoclax (red) or DMSO vehicle (blue). Right panel: summary $BODIPYC_{11}$ staining $697^{WT}$ (black) and $697^{KI}$ (blue) cells in response to 50nM Navitoclax,

200nM Navitoclax or DMSO vehicle normalized to DMSO vehicle. Performed in triplicate, each dot represents a single sample, mean ± SD, two way ANOVA, $^{**}$, *P* = 0.0033

**d,** Left panel: representative histogram of flow cytometric $BODIPYC_{11}$ staining (488nm 530/30) of $REH^{WT}$ (left) and $REH^{Mut1}$ (right) cells in response to 200nM of Venetoclax. Right panel: summary $BODIPYC_{11}$ staining $REH^{WT}$ (green) and $REH^{Mut1}$ (purple) cells in response to Venetoclax normalized to DMSO vehicle. Performed in triplicate, each dot represents a single sample, mean ± SD, two way ANOVA, $^{****}$, *P* <0.0001.

Extended Data Figure 5: Pharmacological inhibition of CREBBP function can sensitize B-ALL to Venetoclax *in-vitro*

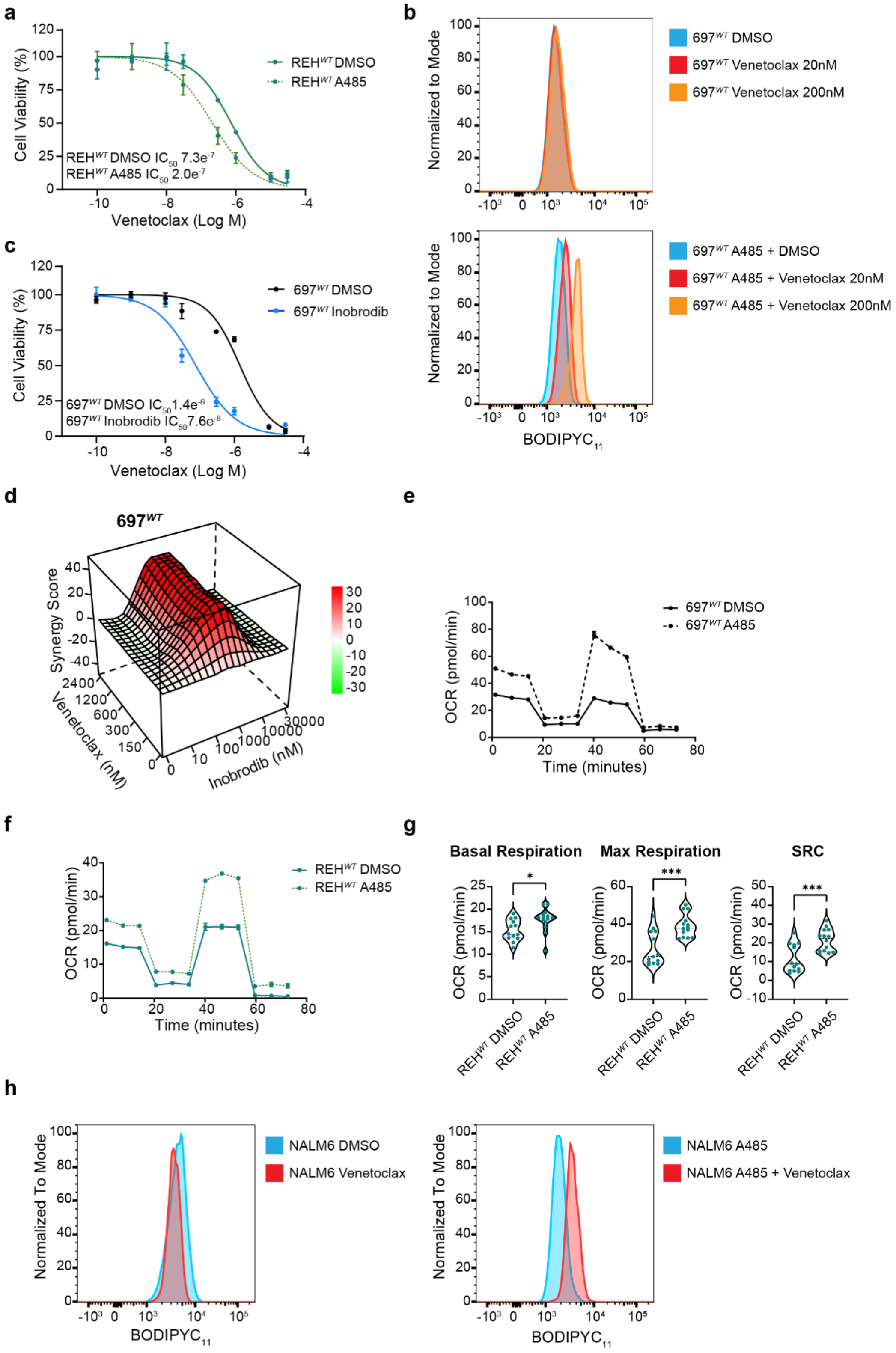

**Extended Data Figure 5: Pharmacological inhibition of CREBBP function can sensitize B-ALL to Venetoclax *in-vitro*.**

**a,** Dose response curve of $REH^{WT}$ (solid line) and $REH^{WT}$ pre-treated with 3 days of A485 (hashed line) to Venetoclax in 72 hour MTS viability assays. Performed in triplicate, mean ± SD.

**b,** Representative histogram of flow cytometric $BODIPYC_{11}$ staining of $697^{WT}$ (top) and $697^{WT}$ pre-treated with A485 (bottom) in response to increasing doses of Venetoclax.

**c,** Dose response curve of $697^{WT}$ (black) and $697^{WT}$ pre-treated with 3 days of Inobrodib (pale blue) to Venetoclax in 72h MTS viability assays. Performed in triplicate, mean ± SD.

**d,** Three-dimensional diffusion plot of ZIP synergy score to combined doses of synchronous Inobrodib and Venetoclax (peak ZIP 37.58). Viability measured by 72h MTS assay.

**e,** Representative mitochondrial oxygen consumption rate (OCR) measured using Seahorse (Agilent) Mitostress test in $697^{WT}$ (solid line) compared to $697^{WT}$ treated with A485 (hashed line). Mean ± SEM.

**f,** Representative mitochondrial oxygen consumption rate (OCR) measured using Seahorse (Agilent) Mitostress test in $REH^{WT}$ (solid line) compared to $REH^{WT}$ treated with A485 (hashed line). Mean ± SEM.

**g,** Summary of basal (left) and maximal (middle) mitochondrial oxygen consumption rate (OCR) and spare respiratory capacity (right) measured using Seahorse (Agilent) Mitostress test in $REH^{WT}$ cells treated with A485 or DMSO vehicle. Each dot represents a single replicate acquired from two separate experiments. Unpaired *t* test, $^{***}$, $P$ >0.0001; $^{*}$, $P$=0.0198.

**h,** Representative histogram of flow cytometric $BODIPYC_{11}$ staining of $NALM6^{WT}$ (left) and $NALM6^{WT}$ pre-treated with A485 (right) in response to Venetoclax.

Extended Data Figure 6: Genetic or pharmacological inhibition of CREBBP sensitizes B-ALL to Venetoclax *in-vivo*.

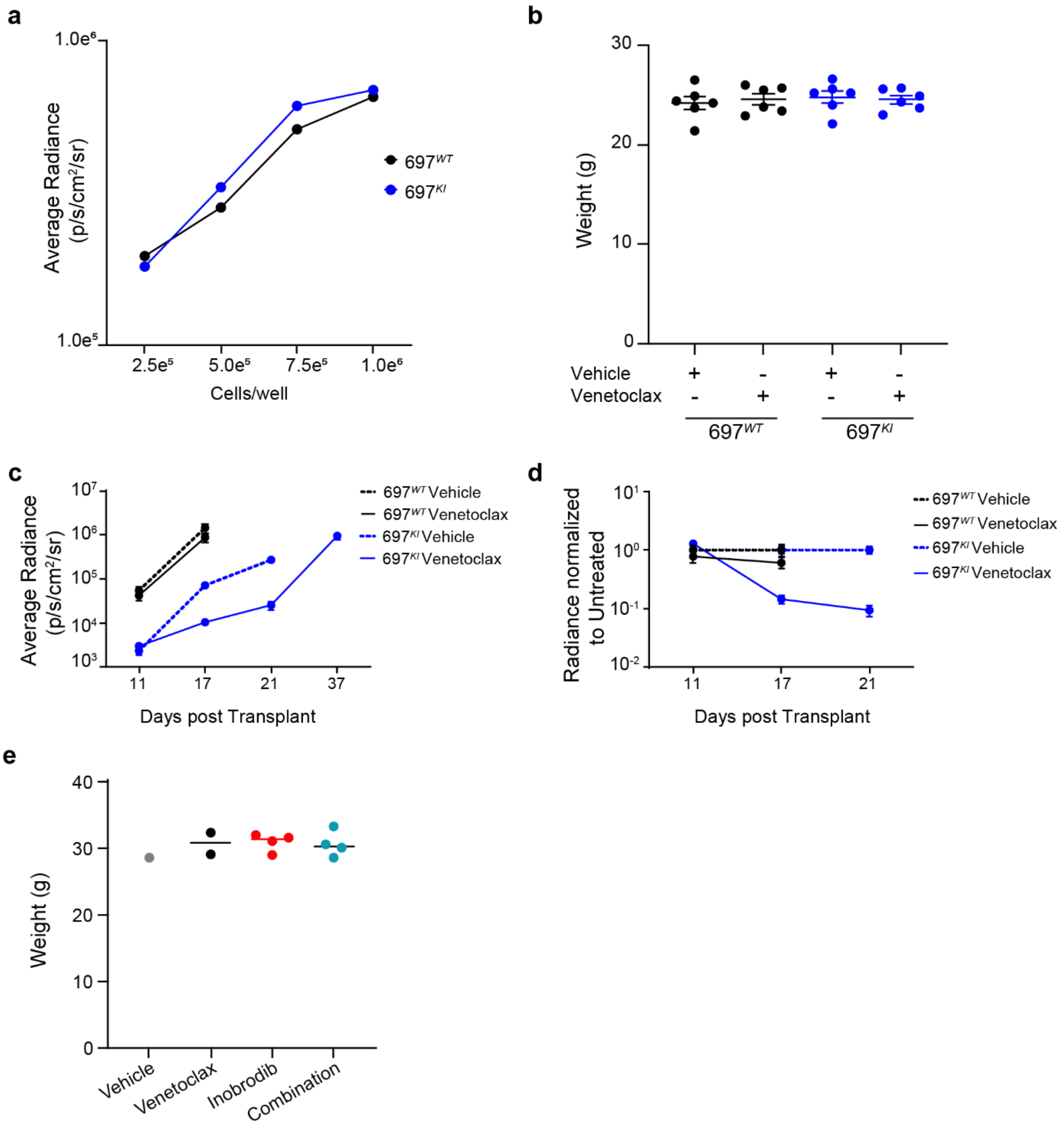


**Extended Data Figure 6: Genetic or pharmacological inhibition of CREBBP sensitizes B-ALL to Venetoclax *in-vivo*.**

**a,** *In-vitro* BLI measurements of a serial dilution of luciferase-expressing $697^{WT}$ vs. $697^{KI}$ B-ALL cells (log p/s/cm$^2$/sr).

**b,** Baseline weights of NSG mice treated in Fig. 7A-F. Mean ± SEM.

**c,** Average BLI radiance of mice engrafted with $697^{WT}$ or $697^{KI}$ cells treated with Venetoclax or vehicle control at days 11, 17 and 21 and 37 ($697^{KI}$ only) of treatment (log p/s/cm$^2$/sr). Mean ± SEM.

**d,** Average BLI radiance of mice engrafted with $697^{WT}$ or $697^{KI}$ cells treated with Venetoclax or vehicle control at days 11, 17 and 21 ($697^{KI}$ only) of treatment (log p/s/cm$^2$/sr) normalized to untreated $697^{KI}$ recipients. Mean ± SEM.

**e,** Baseline weights of NSG mice treated in Fig. 7G. Bar represents median average.
